# DeepCDR: a hybrid graph convolutional network for predicting cancer drug response

**DOI:** 10.1101/2020.07.08.192930

**Authors:** Qiao Liu, Zhiqiang Hu, Rui Jiang, Mu Zhou

## Abstract

**Motivation:** Accurate prediction of cancer drug response (CDR) is challenging due to the uncertainty of drug efficacy and heterogeneity of cancer patients. Strong evidences have implicated the high dependence of CDR on tumor genomic and transcriptomic profiles of individual patients. Precise identification of CDR is crucial in both guiding anti-cancer drug design and understanding cancer biology.

**Results:** In this study, we present DeepCDR which integrates multi-omics profiles of cancer cells and explores intrinsic chemical structures of drugs for predicting cancer drug response. Specifically, DeepCDR is a hybrid graph convolutional network consisting of a uniform graph convolutional network (UGCN) and multiple subnetworks. Unlike prior studies modeling hand-crafted features of drugs, DeepCDR automatically learns the latent representation of topological structures among atoms and bonds of drugs. Extensive experiments showed that DeepCDR outperformed state-of-the-art methods in both classification and regression settings under various data settings. We also evaluated the contribution of different types of omics profiles for assessing drug response. Furthermore, we provided an exploratory strategy for identifying potential cancer-associated genes concerning specific cancer types. Our results highlighted the predictive power of DeepCDR and its potential translational value in guiding disease-specific drug design.

**Availability:** DeepCDR is freely available at https://github.com/kimmo1019/DeepCDR

**Supplementary information:** Supplementary data are available at *Bioinformatics* online.

## 1 Introduction

Designing novel drugs with desired efficacy for cancer patients is of great clinical significance in pharmaceutical industry (Lee *et al.*, 2018). However, the intra- and inter-tumoral heterogeneity results in diverse anti-cancer drug responses among patients (Rubin, 2015; Kohane, 2015), highlighting the complexity of genomics and molecular backgrounds. Recent advances in high-throughput sequencing (HTS) technologies have deepened our understanding of cancer phenotypes from the aggregated amounts of cancer omics profiles (Gagan and Van Allen, 2015). For example, the pharmacogenomics (Daly, 2017; Musa *et al.*, 2017) is evolving rapidly by addressing the interactions between genetic makeup and drug response sensitivity.

Precise identification of cancer drug response (CDR) has become a crucial problem in guiding anti-cancer drug design and understanding cancer biology. Particularly, cancer cell lines (permanently established *in vitro* cell cultures) play an important role in pharmacogenomics research as they reveal the landscape of environment involved in cellular models of cancer (Iorio *et al.*, 2016). Databases such as Cancer Cell Line Encyclopedia (CCLE) (Barretina *et al.*, 2012) provide large-scale cancer profiles including genomic (e.g., genomic mutation), transcriptomic (e.g., gene expression) and epigenomic data (e.g., DNA methylation). Also, the Genomics of Drug Sensitivity in Cancer (GDSC) (Iorio *et al.*, 2016) has been carried out for investigating the drug response to numerous cancer cell lines. For example, the half-maximal inhibitory concentration (IC_50_) is a common indicator reflecting drug response across cancer cell lines. Mining these cancer-associated profiles and their interactions will help characterize cancer molecular signatures with therapeutic impact, leading to accurate anti-cancer drug discovery. However, due to the complexity of omics profiles, the translational potential of identifying molecular signatures that determines drug response has not been fully explored.

So far, a handful of computational models have been proposed for predicting cancer drug response which can be divided into two major categories. The first type is the network-driven methods which analyze the information extracted from drug-drug similarities and cancer cell line similarities. The core idea is to construct a similarity-based model and assign the sensitivity profile of a known drug to a new drug if there are structurally similar. For example, (Zhang *et al.*, 2015) established a dual similarity network based on the gene expression of cancer cell lines and chemical structures of drugs to predict cancer drug response. (Turki and Wei, 2017) proposed a link-filtering algorithm on cancer cell line network followed by a linear regression for predicting the cancer drug response. HNMDRP (Zhang *et al.*, 2018) is a heterogeneous network that integrates multiple networks, including cell line similarity, drug similarity and drug target similarity. An information flow algorithm was proposed for predicting novel cancer drug associations. Notably, network-driven methods tend to show poor scalability and low computational efficiency. Machine learning methods are another type of computational analysis directly exploring profiles from large-scale drugs and cancer cell lines. Typical approaches include logistic regression (Geeleher *et al.*, 2014), Support Vector Machines (SVM) (Dong *et al.*, 2015), random forest (Daemen *et al.*, 2013) and neural networks (Chang *et al.*, 2018; Liu *et al.*, 2019; Sharifi-Noghabi *et al.*, 2019; Manica *et al.*, 2019). Most machine learning methods used single omics data from cancer cell lines, such as genomic mutation or gene expression. For example, CDRscan (Chang *et al.*, 2018) used the molecular fingerprints for drug representation and genomic mutation as cancer cell profile. They were fed to an ensemble CNN model for cancer drug response prediction. tCNNs (Liu *et al.*, 2019) takes SMILES sequence for drug representation and genomic mutation as cancer cell profile, which will be fed to a twin convolutional neural network as inputs. We summarized the major limitations of prior studies as follows.

- Conventional feature extractions are unable to capture intrinsic chemical structures of drugs. For example, engineered features of compounds only consider chemical descriptors and molecular fingerprints (Liu *et al.*, 2018a; Wei *et al.*, 2019; Chang *et al.*, 2018). Although they have been applied to drug discovery and compound similarity search (Cereto-Massagué *et al.*, 2015), such features are sparse and computationally expensive for drug representation. Also, string-based (e.g., SMILES) representation of drugs (Segler *et al.*, 2017; Guimaraes *et al.*, 2017; Popova *et al.*, 2018; Liu *et al.*, 2019) is quite brittle as small changes in the string can lead to completely different molecules (Kusner *et al.*, 2017).
- Despite the emergence of multi-omics profiles, the vast majority of previous studies merely focused on the analysis of single type of omics data, such as genomic or transcriptomic profiles of cancer cells. The synergy of omics profiles and their interplay have not been fully explored. In addition, the epigenomic data (e.g., DNA methylation), proven to be highly related to cancer occurrence (Klutstein *et al.*, 2016), is largely ignored.

Considering the above limitations, we proposed a hybrid graph convolutional network for predicting cancer drug response (Fig 1). DeepCDR consists of a uniform graph convolutional network (UGCN) for drug representation based on the chemical structure of drugs. Additionally, DeepCDR contains several subnetworks for feature extraction of multi-omics profiles from genomics, transcriptomics and epigenomics inputs. The high-level features of drugs and multi-omics data were then concatenated together and fed into a 1-D convolutional neural network (CNN). DeepCDR enables prediction of the IC_50_ sensitivity value of a drug with regard to a cancer cell line in a regression task, or claiming the drug to be sensitive or resistant in a classification task. Conceptually, DeepCDR can be regarded as a multimodal deep learning solution for cancer drug response prediction. We summarized our contributions as follows.

- We proposed a uniform graph convolutional network (UGCN) for novel feature extraction of drugs. Compared to hand-crafted features (e.g., molecular fingerprints) or string-based features (e.g., SMILES), the novel design of UGCN architecture can automatically capture drug structures by considering the interactions among atoms within a compound.
- We discovered that the synergy of multi-omics profiles from cancer cell lines can significantly improve the performance of cancer drug response prediction and epigenomics profiles are particularly helpful according to our analysis.
- We designed extensive experiments to reveal the superiority of our model. DeepCDR achieves state-of-the-art performance in both classification and regression settings, highlighting the strong predictive power of UGCN architecture and multimodal learning strategy.

**Fig. 1.**
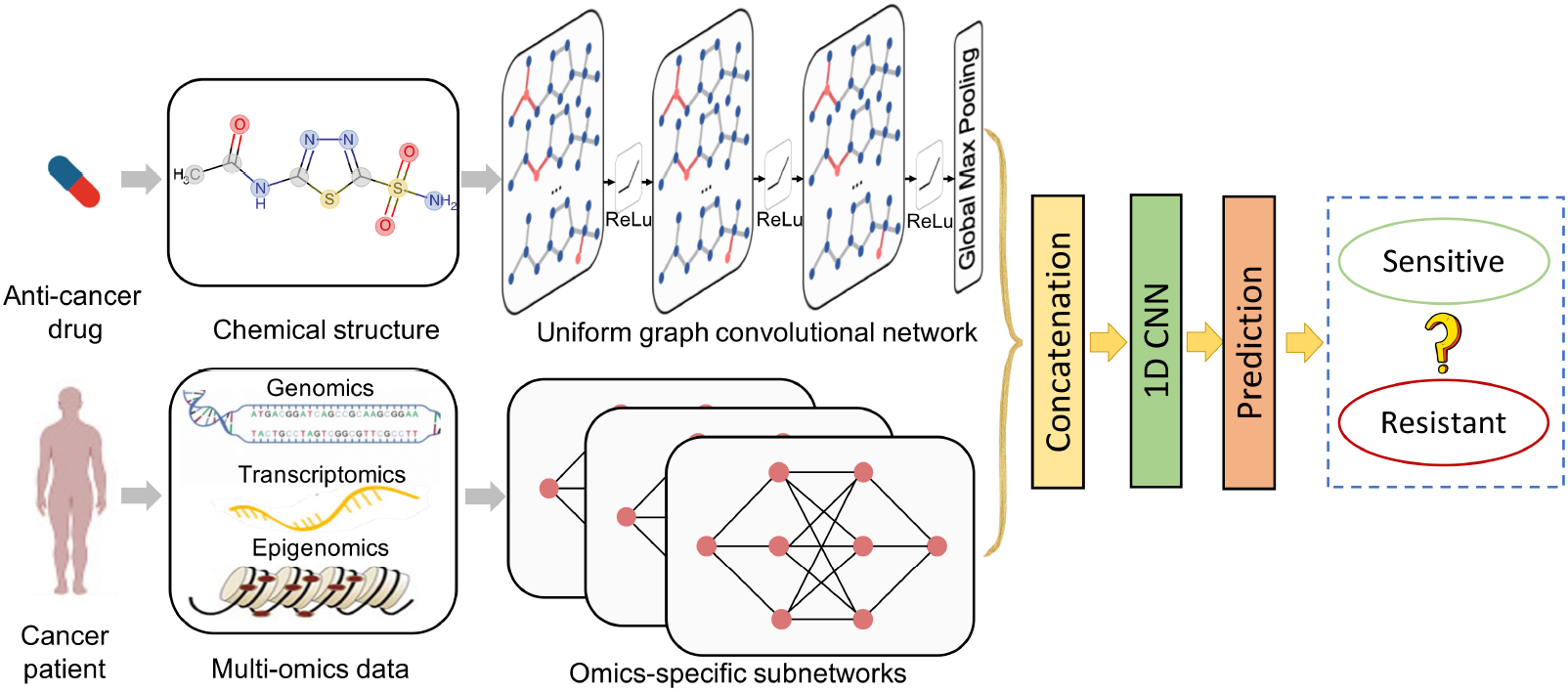
The overview framework of DeepCDR. DeepCDR contains a uniform graph convolutional network (UGCN) and three subnetworks for processing drug structures and cancer cell profiles (genomic mutation, gene expression and DNA methylation data) respectively. DeepCDR takes a pair of drug and cancer cell profiles as inputs and predicts the drug sensitivity (IC_50_) (regression) or claims the drug to be sensitive or resistant (classification). The drug will be represented as a graph based on the chemical structure before transformed into a high-level latent representation by the uniform graph convolutional network (UGCN). Omics featured learned by subnetworks will bes concatenated to the drug feature.

## 2 Methods

### 2.1 Overview of DeepCDR framework

DeepCDR is constructed by a hybrid graph convolutional network for cancer drug response (CDR) prediction, which integrates both drug-level and multi-omics features (Fig 1). The output of DeepCDR is measured by the IC_50_, which denotes the effectiveness of a drug in inhibiting the growth of a specific cancer cell line. For example, small IC_50_ value reveals a high degree of drug efficacy, implying that the drug is sensitive to the corresponding cancer cell line.

DeepCDR consists of a uniform graph convolutional network (UGCN) and several subnetworks for extracting drug and cancer cell line information, respectively (see detailed hyperparameters in Supplementary Table 1-2). On the one hand, the uniform graph convolutional network (UGCN) takes the adjacent information of atoms in a drug into consideration by aggregating the features of neighboring atoms together. On the other hand, the subnetworks extract high-level features of cancer omics profiles from a certain cancer cell line (i.e., genomic data, transcriptomic data and epigenomic data). Then the high-level features of drug and multiple omics data were concatenated and fed to a 1-D convolutional neural network. To alleviate potential overfitting in the training process, we used Batch normalization (Ioffe and Szegedy, 2015) and Dropout (Srivastava *et al.*, 2014) after each convolutional layer. We used Adam as the optimizer for updating the parameters of DeepCDR in the back-propagation process. Similar to (Liu *et al.*, 2018b), the DeepCDR classification model takes a sigmoid layer for prediction and cross-entropy (CE) as loss function, while the DeepCDR regression model directly uses a linear layer without an activation function and takes mean square error (MSE) as loss function.

### 2.2 Drug feature representation

Each drug has its unique chemical structure which can be naturally represented as a graph where the vertices and edges denote chemical atoms and bonds, respectively. Suppose we have *M* drugs in our study, the graph representation of these drugs can be described as 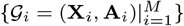 where 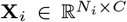 and 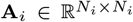 are the feature matrix and adjacent matrix of the *i*^*th*^ drug, respectively. *N*_*i*_ is the number of atoms in the *i*^*th*^ drug and *C* is the number of feature channels. Each row of feature matrix corresponds to the attributes of an atom. Following the description in (Ramsundar *et al.*, 2019), the attributes of each atom in a compound were represented as a 75-dimensional feature vector (*C*=75), including chemical and topological properties such as atom type, degree and hybridization. We downloaded the structural files (.MOL) of all drugs (*M* =223) from PubChem library (Kim *et al.*, 2018) of which the number of atoms *N*_*i*_ varies from 5 to 96.

### 2.3 Uniform graph convolutional network

We seek to achieve graph-level classification as each input of drug represents a unique graph structure while the original graph convolutional network (GCN) (Kipf and Welling, 2017) aims at node classification within a single graph. To address this issue, we extended the original GCN architecture and presented a uniform graph convolutional network (UGCN) for processing drugs with variable sizes and structures. The core idea of UGCN is to introduce an additional complementary graph to the original graph of each drug to ensure the consistent size of feature matrix and adjacent matrix. Given the original graph representation of *M* drugs 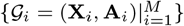, the complementary graphs of drugs can be represented as 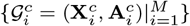, where 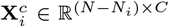 and 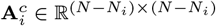. *N* is a fixed number which is set to 100. Thus the consistent representation of a drug is designed as follows:

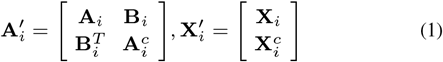

where 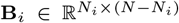 is a conjunction matrix which represents the connection between the *i*^*th*^ original graph and complementary graph. 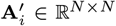 and 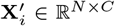 are the consistent adjacent matrix and feature matrix. The uniform graph convolutional network (UGCN) applied to *i*^*th*^ drug is defined as 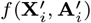 with a layer-wise operation as

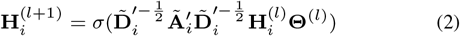

where 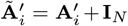 is the adjacent matrix with added self-connections, 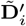 is the degree matrix of 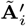 which 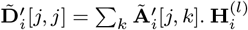 and **Θ**^(*l*)^ are the convolved signal and filter parameters of the *l*^*th*^ layer. *σ*(·) is the activation function, which is set to *ReLu*(·) = *max*(0, ·). We further denote the first *N*_*i*_ rows of 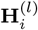 as 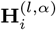 and the remaining (*N − N*_*i*_) rows as 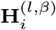. For the first graph layer where *l* = 0, we initialized the first layer as 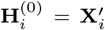 and substituted formula (1) into (2), we can derive the propagation rule of first layer of UGCN as the following:

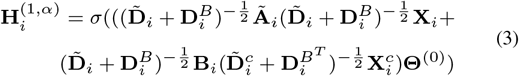

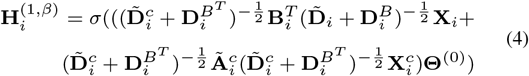

where 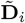 and 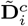 are the degree matrix of 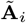 and 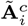 and 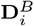 and 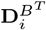 are two diagonal matrix for describing row sum and column sum of **B**_*i*_. 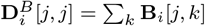 and 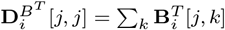.

With mathematical induction, it can be inferred that the general layer-wise propagation rule of UGCN can be represented by the following two equations:

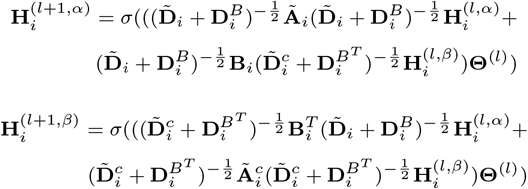

We further consider a special case where the complementary graphs have no connection to the original graphs (**B**_*i*_ = **0**), so the layer-wise propagation rule of UGCN will be simplified as

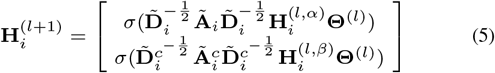

Overall we showed that the convolution on the original graph and the corresponding complementary graph is independent in UGCN given the conjunction matrix **B**_*i*_ = **0**. At last, we applied a global max pooling (GMP) over the graph nodes in 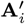 to ensure that drugs with different size will be embedded into a fixed dimensional vector (defulat dimension: 100). In our study, we set **B**_*i*_ = **0**, 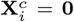 and *l* = 3 as the default settings for initializing DeepCDR. We also explored another initailization strategy in the discussion section.

### 2.4 Omics-specific subnetworks

We designed omics-specific subnetworks to integrate the information of multi-omics profiles. We used the late-integration fashion in which each subnetwork will first learn a representation of a specific omics data in a latent space and then be concatenated together. The three subnetworks can be represented as {***y***_*g*_=***f***_*g*_ (***x***_*g*_), ***y***_*t*_=***f***_*t*_(***x***_*t*_), ***y***_*e*_=***f***_*e*_(***x***_*e*_)} for processing **g**enomic, **t**ranscriptomic and **e**pigenomic data per sample, respectively. Similar to (Chang *et al.*, 2018), we used a 1D convolutional network for processing genomic data as the mutation positions are distributed linearly along the chromosome. For transcriptomic and epigenomic data, we directly used fully-connected networks for feature representation. (see detailed hyperparameters of subnetworks in Supplementary Table 2).

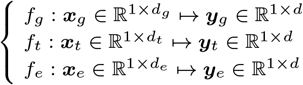

The dimension of latent space *d* is set to 100 in our experiments by detault.

### 2.5 Data preparation

We integrated three public databases in our study including Genomics of Drug Sensitivity in Cancer (Iorio *et al.*, 2016) (GDSC), Cancer Cell Line Encyclopedia (Barretina *et al.*, 2012) (CCLE) and TCGA patient data (Weinstein *et al.*, 2013). GDSC database provides IC_50_ values for a large-scale drug screening data, of which each IC_50_ value corresponds to a drug and a cancer cell line interaction pair. CCLE database provides genomic, transcriptomic and epigenomic profiles for more than a thousand cancer cell lines. For the three omics data, we focused on genomic mutation data, gene expression data and DNA methylation data, which can be easily accessed and downloaded using DeMap portal (https://depmap.org). TCGA patient data provide both genetic profiles of patients and clinic annotation after drug treatment. We used TCGA dataset for an external validation.

We downloaded IC_50_ values (natural log-transformed) across hundreds of drugs and cancer cell lines from GDSC database as the ground truth of drug sensitivity profiles for measuring cancer drug response. We excluded drug samples without PubChem ID in GDSC database and removed cancer cell lines in which any type of omics data was missing. Note that several drugs with different GDSC ids may share the same PubChem ids due to the different screening condition. We treated them as individual drugs in our study. We finally collected a dataset containing 107446 instances across 561 cancer cell lines and 238 drugs. Considering all the 561 × 238 = 133518 drug and cell line interaction pairs, approximately 19.5% (26072) of the IC_50_ values were missing. The corresponding drug and cancer cell line datasets used in this study are summarized in Supplementary Table 3-4. Each instance corresponds to a drug and cancer cell line interaction pair. Each cell line was annotated with a cancer type defined in The Cancer Genome Atlas (TCGA) studies and we only considered TCGA cancer types suggested by (Chang *et al.*, 2018) in the downstream analysis.

For multi-omics profiles of cancer cell lines, we only consider data related to 697 genes from COSMIC Cancer Gene Census (https://cancer.sanger.ac.uk/census). For genomic mutation data, 34673 unique mutation positions including SNPs and Indels within the above genes were collected. The genomic mutation of each cancer cell line was represented as a binary feature vector in which ‘1’ denotes a mutated position and ‘0’ denotes a non-mutated position (*d*_*g*_ =34673). For gene expression data, the TPM value of gene expression was log_2_ transformed and quantile normalized. Then the gene expression of each cell line can be represented as a 697-dimensional feature vector (*d*_*t*_=697). The DNA methylation data was directly obtained from the processed Bisulfite sequencing data of promoter 1kb upstream TSS region. Then we applied a median value interpolation to the data as there were a minority of missing values. The methylation of each cell line is finally represented by a 808-dimensional feature vector (*d*_*e*_=808). The three types of omics data were finally transformed into a latent space where the embedded dimension was fixed to 100 (*d*=100).

For TCGA patient data, we chose the patients with cervical squamous cell carcinoma and endocervical adenocarcinoma (CESC) disease with two criterions. 1) The gene mutation, gene expression and DNA methylation data are available. 2) The clinic annotation of drug response was also available. We finally created an external data source with 54 records across 31 patients and 12 drugs. The genetic profiles were preprocessed the same way as cell line data and we took records with “Complete Response” clinic annotation as positive examples (see details in Supplementary Table 5).

### 2.6 Baseline methods

The following competing methods were considered. The best or default parameters of each method were used for model comparison.

- **Ridge Regression** is a linear regression model with *L*_2_ penalty. We first concatenated genomic mutation features and molecular fingerprints of drugs together and then fed to ridge regression model. The ridge regression model was implemented using sklearn library (Pedregosa *et al.*, 2011). Basically, we found no significant changes in results as we tried different settings of the *L*_2_ penalty coefficient from {0.1,0.5,1.0,5.0}. We finally chose the default coefficient parameter 1.0 provided by sklearn library in the comparing experiments.
- **Random Forest** is a tree-based regressor in which the input is the same as the ridge regression model. The random forest was also implemented with sklearn library (Pedregosa *et al.*, 2011). We set the number of trees in the forest from {10,100,200,500} and chose the best parameter in the comparing experiments.
- **CDRscan** (Chang *et al.*, 2018) applies an ensemble convolutional neural networks (CNNs) model for predicting cancer drug response using molecular fingerprints of drugs and genomic mutation data of cancer cell line.
- **tCNNs** (Liu *et al.*, 2019) applies an convolutional neural network (CNN) for predicting cancer drug response using SMILES sequences of drugs and genomic mutation data of cancer cell line.SMILES sequences of drugs will first be encoded into one-hot representation and fed to the neural network.
- **MOLI** (Sharifi-Noghabi *et al.*, 2019) is one of the few studies that considers multi-omics profiles (genomic mutation, copy number and gene expression) with encoder neural networks. Specifically, MOLI is a drug-specific model where each model is trained for a specific drug.

### 2.7 Model evaluation

In the regression experiments for predicting natural log-transformed IC_50_ values given the profiles of drugs and cancer cell lines, we used three common metrics for measuring the statistical correlation between observed values and predicted IC_50_ values, including Pearson’s correlation coefficient (PCC), Spearman’s correlation coefficient (SCC) and root mean squared error (RMSE). PCC measures the linear correlation between observed and predicted IC_50_ values while SCC is a nonparametric measure of rank correlation of observed and predicted IC_50_ values. RMSE directly measures the difference of observed and predicted IC_50_ values.

For classification experiments, we chose the area under the receiver operating characteristic curve (AUC) and area under the precision-recall curve (auPR) as the two commonly used measurements of a classifier.

To comprehensively evaluate the performance of our model DeepCDR, we demonstrated results under various data settings. We breifly summarized these different data settings in the following:

- **Rediscovering Known Cancer Drug Responses**. Based on the known drug-cell line interactions across 561 cancer cell lines and 238 drugs, we randomly selected 80% of instances of each TCGA cancer type as the training set and the remaining 20% of the instances as the testing set for model evaluation. The five-fold cross-validation was conducted.
- **Predicting Unknown Cancer Drug Responses**. We trained DeepCDR model with all the known drug-cell line interaction pairs and predicted the missing pairs in GDSC database (approximately 19.5% of all pairs across 561 cancer cell lines and 238 drugs).
- **Blind Test for Both Drugs and Cell Lines**. In order to evaluate the predictive power of DeepCDR when given a new drug or new cell line that is not included in the training data. We randomly split the data into 80% training set and 20% test set on the cell line or drug level. The five-fold cross-validation using leave drug out and leave cell line out strategy was conducted.
- **External Validation with Patient Data**. To evaluate whether DeepCDR trained with *in vitro* cell line data can be generalized to *in vivo* patient data. We trained DeepCDR classification model with cell line data and tested on TCGA patient data.

## 3 Results

### 3.1 DeepCDR recovers continuous degree of drug sensitivity

We first designed a series of experiments to see whether DeepCDR can help recover continuous degree of drug sensitivity. For this objective, we created datasets of the drug and cancer cell lines profiles from GDSC (Iorio *et al.*, 2016) and CCLE (Barretina *et al.*, 2012) database, respectively. We then evaluated the regression performance of DeepCDR and five comparing methods based on the observed IC_50_ values and predicted IC_50_ values. Three common regression evaluation metrics, including Pearson’s correlation coefficient (PCC), Spearman’s correlation coefficient (SCC) and root mean square error (RMSE), were considered.

First, we evaluated the ability of DeepCDR and competing methods by rediscoverying cancer drug response across multiple drugs and cell lines. We observed that DeepCDR demonstrated superior predictive performance of drug response in the regression experiments by achieving the highest Pearson’s correlation and Spearman’s correlation and lowest RMSE comparing to five competing methods (Table 1). Generally, deep neural network models significantly outperformed other baselines as linear or tree-based model may not well capture the intrinstic structural information within drugs. Among the four deep learning models, DeepCDR outperforms three other deep learning methods with a relatively large margin by achieving a Pearson’s correlation of 0.923 as compared to 0.813 of MOLI, 0.871 of CDRscan and 0.885 of tCNNs. This conclusion is also consistent considering other metrics such as Spearman’s correlation and root mean square error (RMSE). Additionally, we also showed the model variance by indepedent training for five times.

**Table 1.** Regression experiments of IC_50_ values with DeepCDR and five comparing methods. Three different measurements, including Pearson’s correlation, Spearman’s correlation and root mean square error (RMSE), were illustrated. We trained neural network based models from scratch for five times and the standard deviations of each method were also calclated for evaluating the model robustness. DeepCDR demonstrates a consistant highest performance in all measurements comparing to other methods

| Methods | Pearson's correlation | Spearman's correlation | RMSE |
| --- | --- | --- | --- |
| Ridge Regression | 0.780 | 0.731 | 2.368 |
| Random Forest | 0.809 | 0.767 | 2.270 |
| MOLI | $0.813 \pm 0.007$ | $0.782 \pm 0.005$ | $2.282 \pm 0.008$ |
| CDRscan | $0.871 \pm 0.004$ | $0.852 \pm 0.003$ | $1.982 \pm 0.005$ |
| tCNNS | $0.885 \pm 0.008$ | $0.862 \pm 0.006$ | $1.782 \pm 0.006$ |
| **DeepCDR** | **$0.923 \pm 0.006$** | **$0.903 \pm 0.004$** | **$1.058 \pm 0.006$** |

Next, we illustrated several prediction cases across multiple TCGA cancer types or different drug compounds. Among the 30 different TCGA cancer types, DeepCDR reveals a consistently high performance by achieving a Pearson’s correlation ranging from 0.889 to 0.941. The best prediction case in mutiple myeloma cancer type and the worst prediction case in acute myelold leukemia cancer type were shown in Fig 2A and Fig 2B, respectively. In the perspective of drug, we also evaluated the regression performance with respect to a specific drug. We observed that the DeepCDR illustrates a relatively more dynamic regression performance by achieving a Pearson’s correlation ranging from 0.328 to 0.938 (Fig 2C-2D), which may due to the drug similarity diversity. We then validated this conclusion by measuring the drug similarity among training set and found that Belinostat has a significantly higher drug similarity score compared to Pazopanib (Supplementary Fig.1, *p*-value=2.38×10^−37^). The distribution of correlation across TCGA cancer types and drugs were provided in Supplementary Fig.2.

**Fig. 2.**
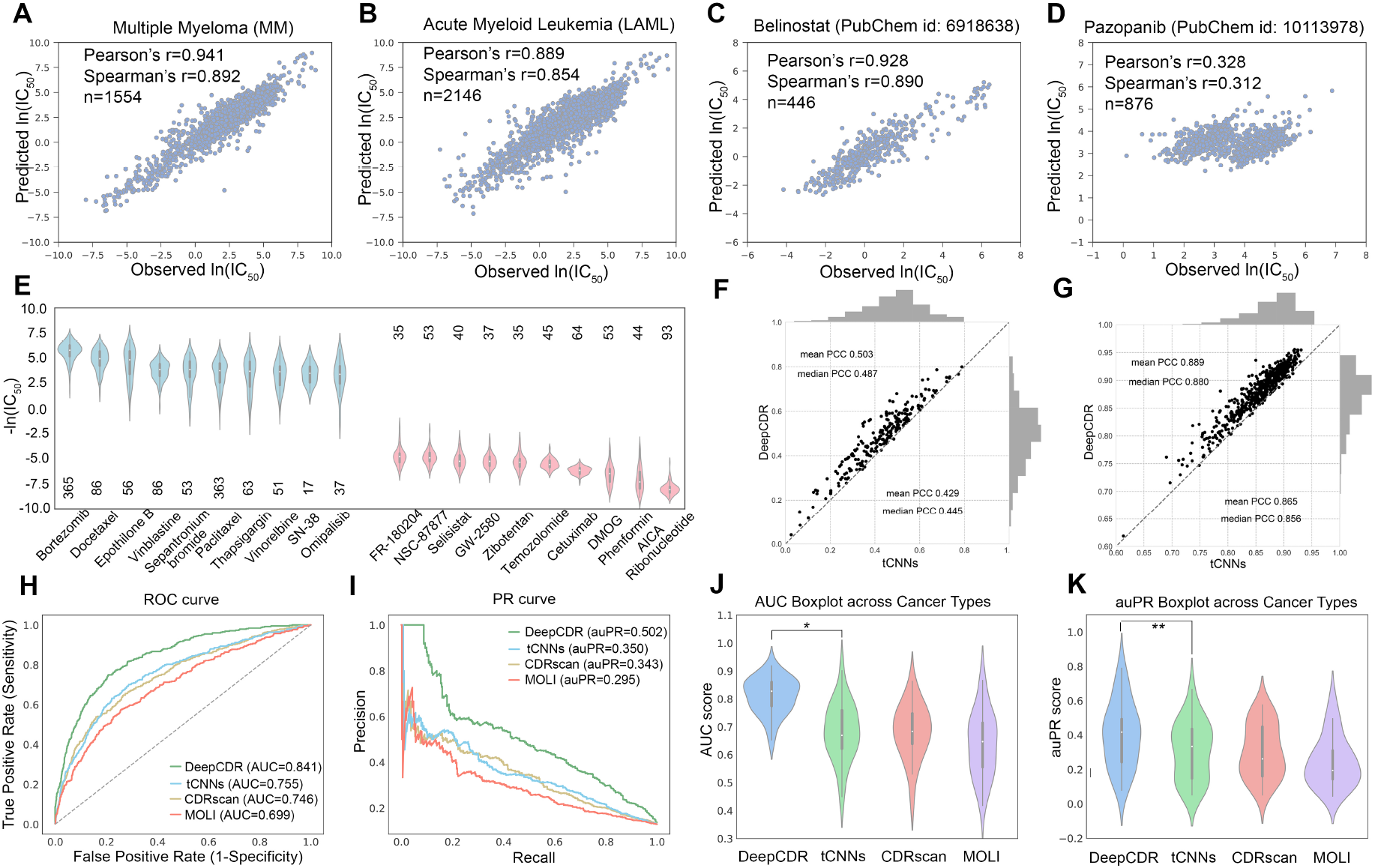
Performance of DeepCDR in cancer drug response prediction under different experiment settings. (A) and (B) highlighted the scatter plots in two TCGA cancer types with the best (MM) and worst (LAML) performance. (C) and (D) showed the scatter plots in two drugs with the best (Belinostat) and worst (Pazopanib) performance. (E) The predicted IC_50_ values of missing data in GDSC database grouped by drugs. Drugs were sort according to the average predicted IC_50_ value in missing cell lines. The number of missing cell lines for each drug is also denoted below/above the violin plot. Each violin plot corresponds to a specifc drug response in all missing cell lines. The blue and red violin plots denote the top-10 drugs with the highest and the lowest efficacy.(F) The performance of DeepCDR and tCNNs in blind test for drugs. The x-axis and y-axis of each dot represent the Pearson’s correlation of tCNNs and DeepCDR, respectively. The dot fallen into the left upper side denotes the case where DeepCDR outperforms tCNNs. (G) The performance of DeepCDR and tCNNs in blind test for cell lines. (H) and (I) show the receiver operating characteristic (ROC) and precision-recall (PR) curve of the four comparing methods, respectively. (J) and (K) show the violin plots of the area under ROC curve (AUC) and area under PR curve (auPRs) across TCGA cancer types. Note that each dot within a violin plot represents the average AUC or auPR score within one TCGA cancer type. Additionally, one-sided Mann-Whitney U tests between DeepCDR and tCNNs were conducted. *p-value=1.01×10^−5^, **p-value=0.062.

Next, we applied DeepCDR to predicting unknown cancer drug responses in the GDSC database. Towards this goal, DeepCDR was trained on all known drug cell line interaction pairs across 561 cell lines and 238 drugs, then it was used for predicting the missing pairs in GDSC database (approximately 19.5% of all pairs). Fig 2E illustrates the distributions of predicted IC_50_ values in GDSC database grouped by drugs. Note that drugs were sorted by the median predicted IC_50_ value across all missing cell lines. We provided the predicted IC_50_ values of top-10 drugs and related cancer cell lines in Supplementary Table 6. Interestingly, Bortezomib was the drug with highest efficacy in our prediction which has been proved to be a proteasome inhibitor that has activity in multiple cancer cell lines (Richardson *et al.*, 2003). Specifically, the predicted IC_50_ of Bortezomib with a oesophagus cell line KYSE-510 is 7.45 × 10^−5^ which implies a strong therapeutic effect. This prediction was supported by the findings in (Lioni *et al.*, 2008), which highlighted the robust activity of Bortezomib in esophageal squamous cells. Phenformin and AICA ribonucleotide are predicted to have the lowest efficacy. The former was used for treating type 2 diabetes mellitus by inhibiting complex I (Marchetti *et al.*, 1987). The latter is capable of stimulating AMP-dependent protein kinase (AMPK) activity (Corton *et al.*, 1995). The anti-cancer of the two drugs might be not their main function but the side effect.

At last, we designed a series of blind tests for both drugs and cell lines. The task becomes much more challenging as the drugs or cell lines in the test data were unseen during the training process. The drug sensitivity data from GDSC database were split into training and test sets on the drug or cell line level. We compared our model DeepCDR to the best baseline model tCNNs in the previous experiments. In the blind test for drugs, the performance of both methods largely decreased compared to previous experiments. However, DeepCDR still achieves an average Pearson’s correlation of 0.503, compared to 0.429 of tCNNs (Fig 2F, *p*-value<1.51×10^−6^, Supplementary Table 7). In the blind test for cell lines, DeepCDR again outperforms tCNNs by a quite large margin by achieving an average Pearson’s correlation of 0.889, compared to 0.865 of tCNNs (Fig 2G, *p*-value<2.2×10^−16^, Supplementary Table 8).

### 3.2 DeepCDR predicts binary drug sensitivity status

In this section, we binarized IC_50_ according to the threshold of each drug provided by (Iorio *et al.*, 2016). After filtering drug samples without a binary threshold, we collected a dataset with 7488 positive instances in which drugs are sensitive to the corresponding cancer cell line and 52210 negative instances where drugs are resistant to cancer cell lines. Similar to the regression experiment settings, we first compared DeepCDR to three other neural network models by rediscovering the cancer drug response status. Despite of the unbalanced dataset (around 1:7), DeepCDR outperforms three other methods by a large margin by achieving a significantly higher AUC and auPR score of 0.841 and 0.502 (Fig 2H-I), reaffirming the advance of DeepCDR in capturing the interaction information of drug and cancer cells. As seen in Fig 2J-K, we grouped the test instances (each instance denotes a drug and cancer cell line pair) according to the TCGA cancer types, then we calculated the AUCs and auPRs of the two methods under different cancer type groups. We observed that DeepCDR achieves higher AUC score and auPR score than tCNNs with respect to every TCGA cancer type. In the blind test for both drugs and cell lines, DeepCDR achieves a consistently better performance than the best baseline tCNNs with average AUC of 0.737 (Supplementary Fig 3-4). Besides, statistical hypothesis tests, including binomial exact test and Mann-Whitney U test, were additionally conducted in both blind test experiments for drugs and cell lines (Supplementary Table 9-10).

Last, we introduced TCGA patient data as an external validation. We trained DeepCDR model on *in vitro* cell line data described above, and tested on *in vivo* patient data. Note that the external dataset even contains more than 40% of drugs that were not included in the cell line data. DeepCDR still achieves an performance with AUC 0.688, compared to 0.618 of tCNNs (Supplementary Fig 5).

### 3.3 Model ablation analysis

Since most early studies only considered single type of omics data, it is necessary for us to evaluate the contribution of different types of omics data. For each type of omics data, we discarded other types of omics data and trained DeepCDR regression model from scratch for model ablation analysis. When using single omic data, the Pearson’s correlation of DeepCDR ranges from 0.878 to 0.890, indicating the usefulness of all individual omics profiles (Table 2). In particular, the epigenomics data (DNA methylation) contributes the most among different omics profiles. Notably, DeepCDR still achieved a higher Pearson’s correlation than tCNNs even only genomic data was used in both methods (0.889 vs 0.885). Furthermore, to verify the effectiveness of graph convolution, we first eliminated the adjacent information of atoms within drugs by setting adjacent matrices to identity matrices 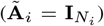. Then the DeepCDR model without adjacent information achieved a reasonable Pearson’s correlation of 0.886. We concluded that the regression performance can be significantly boosted with the powerful representation inferred from adjacent information by the proposed UGCN architecture (0.923 vs 0.886).

**Table 2.** Model ablation studies with different experimental settings. We showed both the contribution of each omic profile and the contribution of graph convolution module

| Experimental setting | Pearson's correlation |
| --- | --- |
| Single genomics | 0.889 |
| Single transcriptomics | 0.878 |
| Single epigenomics | 0.890 |
| Multi-omics without adjacent info | 0.886 |
| Multi-omics with adjacent info | **0.923** |

### 3.4 DeepCDR helps prioritize cancer-associated genes

To deepen the understanding of biological knowledge revealed by DeepCDR, we further proposed an exploratory strategy for prioritizing cancer-associated genes given an input drug and cancer cell line pair, where we prioritized the involved genes by assigning each gene with an associated score. In detail, to obtain the associated scores for 697 genes involved in COSMIC Cancer Gene Census, we considered the absolute gradient of the predicted outcome from DeepCDR regression model with respect to the each gene’s expression. We highlighted several cases where the drugs were shown sensitive to the corresponding cancer cell lines (Table 3). Importantly, we found that many top-ranked genes have been verified to be associated with cancers by existing literature. For example, Erlotinib and Lapatinib, two known drugs for treating lung cancer, have been proven to be EGFR inhibitors (Sayar *et al.*, 2014), EGFR gene ranks first and fourth from DeepCDR prioritizer in A3/KAW and BT-474 cell lines, respectively. Also, Nilotinib is a potential drug treatment for chronic myelogenous leukemia (Kantarjian *et al.*, 2011). Interestingly, in our predictive task in a BHT-101 cell line, ABI1 ranked as the second cancer-associated gene, which has been previously proved to have specific expression patterns in leukemia cell lines (Shibuya *et al.*, 2001). Taken together, these evidences support that DeepDCR could reveal potential therapeutic targets for anti-cancer drugs and help discover hypothetical cancer-associated genes for additional clinical testing.

**Table 3.** Top-5 cancer-associated genes prioritized by DeepCDR. We proposed a simple gradient-based strategy for prioritizing all the genes when making a prediction of a specific drug and cancer cell line pair. Many top-ranked genes have been verified to be highly associated with cancers by existing literatures

| Drug | Cell line | TCGA type | Top-5 cancer-associated genes | ln(IC<sub>50</sub>) |  |
| --- | --- | --- | --- | --- | --- |
|  |  |  |  | Observed | Predicted |
| Erlotinib | A3/KAW | DLBC | EGFR, ALK, BCL10, CREB3L1, STAG1 | 1.110 | 1.206 |
| Lapatinib | BT-474 | BRCA | ERBB2, MDS2, FOXL2, EGFR, MNX1 | -1.028 | -0.879 |
| Bleomycin | A-375 | SKCM | ACKR3, ASXL 1, MTCP1, FOXL2, SALL4 | -1.514 | -1.428 |
| Nilotinib | BHT-101 | THCA | CBLC, ABI1, POU5F1, KLF4, ZNF198 | -0.630 | -0.714 |
| Salubrinal | SUP-B15 | ALL | JAK3, EIF1AX, NUMA1, PRDM1, IL21R | 1.781 | 1.471 |

## 4 Discussion

In this study, we have proposed DeepCDR as an end-to-end deep learning model for precise anti-cancer drug response prediction. We found that graph convolutional networks (GCNs) were extremely helpful for capturing structural information of drugs according to our anaylysis. To the best of our knowledge, DeepCDR is the first work to apply GCN in cancer drug response (CDR) problem. In addition, we demonstrated that the combination of multi-omics profiles and intrinsic graph-based representation of drugs are appealing for assessing drug response sensitivity. Extensive experiments highlighted the predictive power of DeepCDR and its potential translational value in guiding disease-specific drug design.

We provide two future directions for improving our method. 1) The proposed uniform graph convolutinal network (UGCN) can be ultilized for data augmentation when training instances were not adequate enough or extremely unbalanced by randomly sampling multiple complementary graphs for each drug. In the classification experiment where the training data is unbalanced, if we randomlized the feature matrix and gave random connections of complementary graphs and augmented the positive training instances by five times, the average AUC can be further improved by 0.8%. Augmentation with UGCN can potentially further improve prediciton performance. 2) DeepCDR can be leveraged in combination with molecule generation tasks. Current molecule generation models based on RNN language models (Segler *et al.*, 2017), generative adversarial networks (GANs) (Guimaraes *et al.*, 2017) and deep reinforcement learning (Popova *et al.*, 2018) focused on generating general compounds and ignored profiles of targeted cancer cell. Methods focused on cancer-specific or disease-specific novel drug design can be proposed by using CDR predicted by DeepCDR as a prior knowledge or a reward score for guiding molecule generation.

To sum up, we introduced DeepCDR that can be served as an application for exploring drug sensitivity with large-scale cancer multi-omics profiles. DeepCDR outperforms multiple baselines and our analysis illustrates how our method can help prioritize therapeutic targets for anti-cancer drug discovery. In future work, we plan to expand data inclusion for a large-scale omics data profiled both before and after treatment to assess how their molecular profiles respond to perturbation by the testing drugs.

## Acknowledgements

We thank Fengling Chen and Zhana Duren for their helpful discussion.

