## Supplementary Materials for "DeepCDR: a hybrid graph convolutional network for predicting cancer drug response"

Qiao Liu[1], Zhiqiang Hu[2], Rui Jiang[1,*] and Mu Zhou[3,*]
[1] Bioinformatics Division, BNRIST, Tsinghua University, Beijing 100084, China; Department of Automation, Tsinghua University, Beijing 100084, China.
[2] SenseTime Research, Shanghai, China.
[3] SenseBrain Research, CA, USA.

### Contents

**Supplementary Figures** .......... 1

**Supplementary Tables** .......... 7

### Supplementary Figures

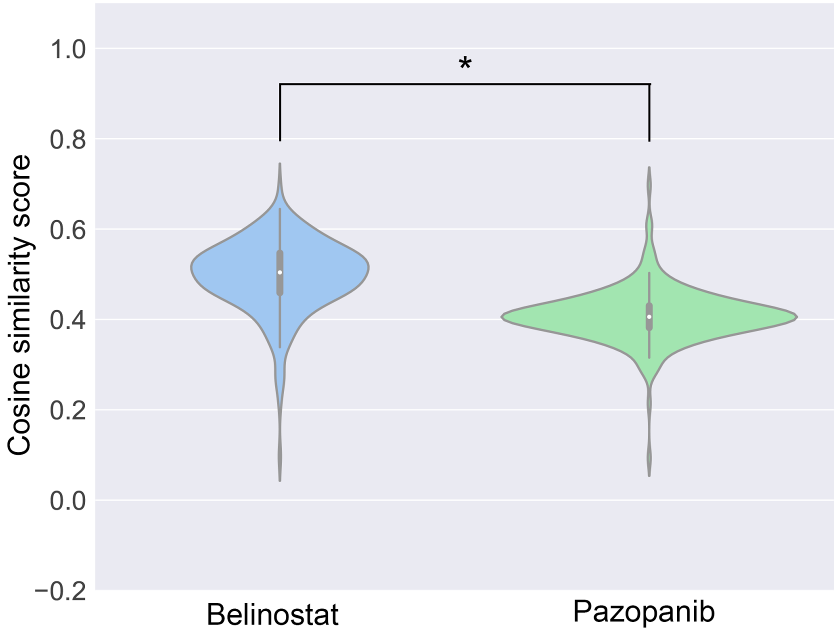


**Supplementary Figure 1.** For the given two drugs Belinostat and Pazopanib, we calculated the cosine similarity scores between itself and other drugs in the training set based on the 3072-dimensional features from molecular fingerprints. We can see that the drug with best regression performance has a significantly higher similarity scores than the drug with the worst regression performance (one-sided Mann-Whitney U test, *p*-value=2.38e-37).

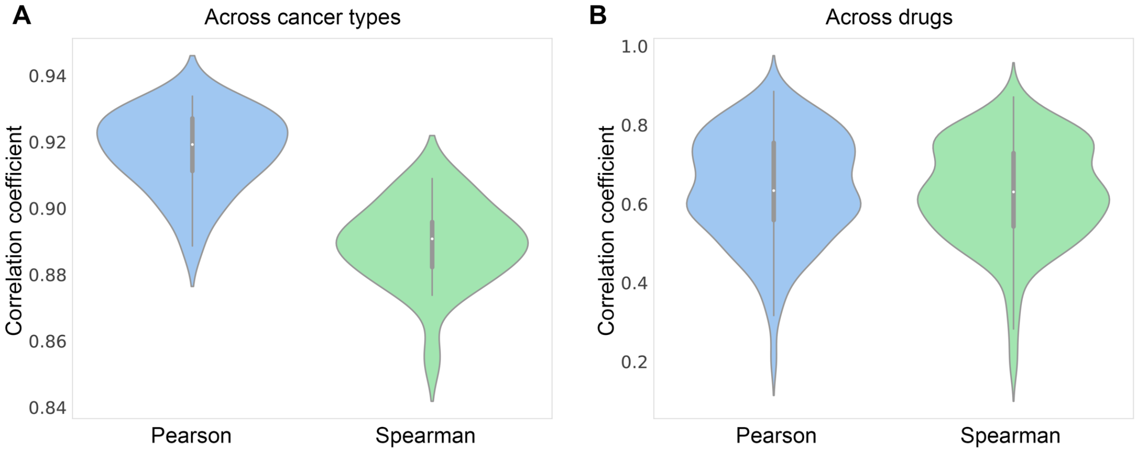


**Supplementary Figure 2.** The performance of DeepCDR across TCGA cancer types and across drugs, respectively. Pearson correlation coefficient and Spearman correlation coefficient were shown.

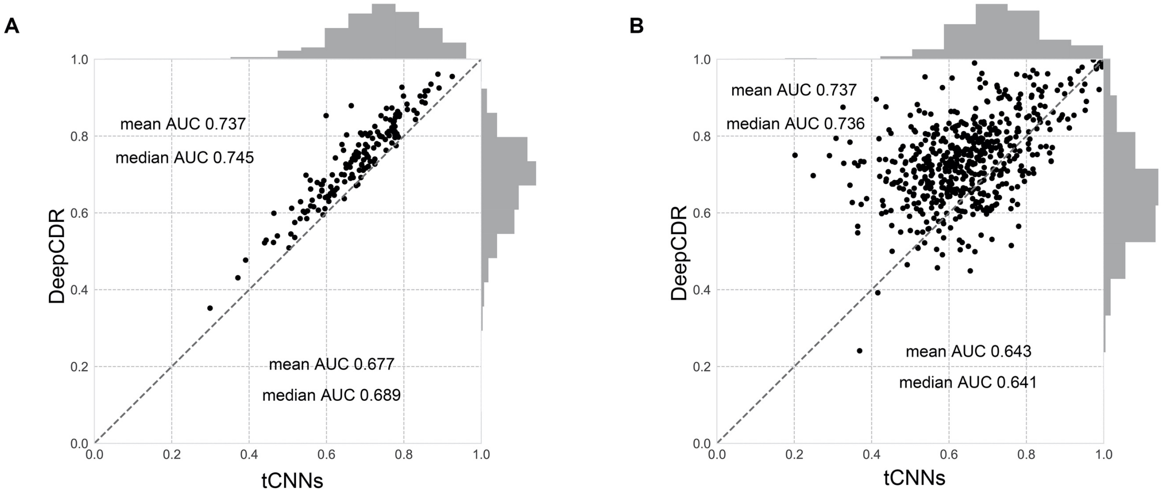


**Supplementary Figure 3.** The performance of DeepCDR and tCNNs in blind test for drug and cell line, respectively. The x-axis and y-axis of each dot represent the AUC of tCNNs and DeepCDR with respect to a specific drug or cell line.

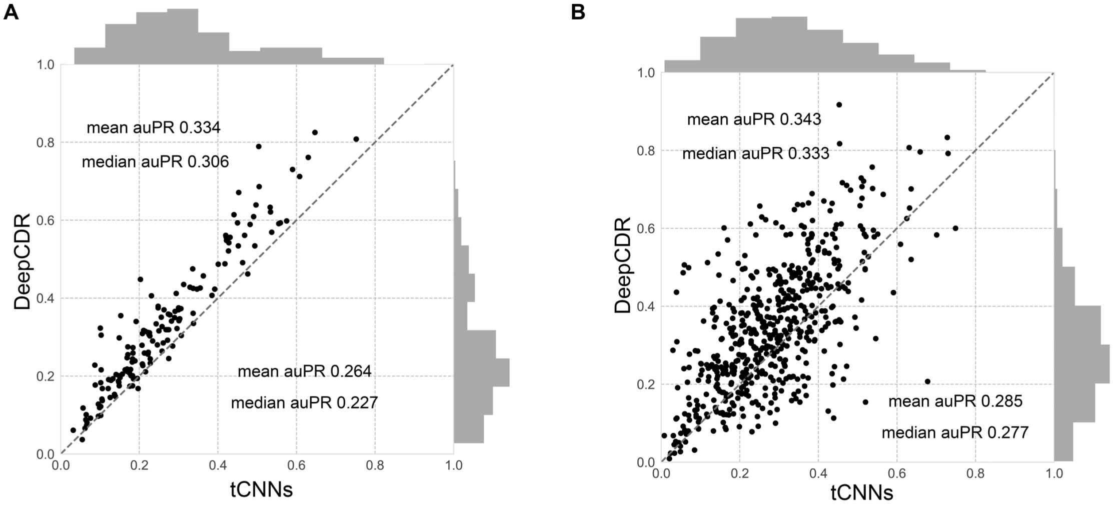


**Supplementary Figure 4.** The performance of DeepCDR and tCNNs in blind test for drug and cell line, respectively. The x-axis and y-axis of each dot represent the auPR of tCNNs and DeepCDR with respect to a specific drug or cell line.

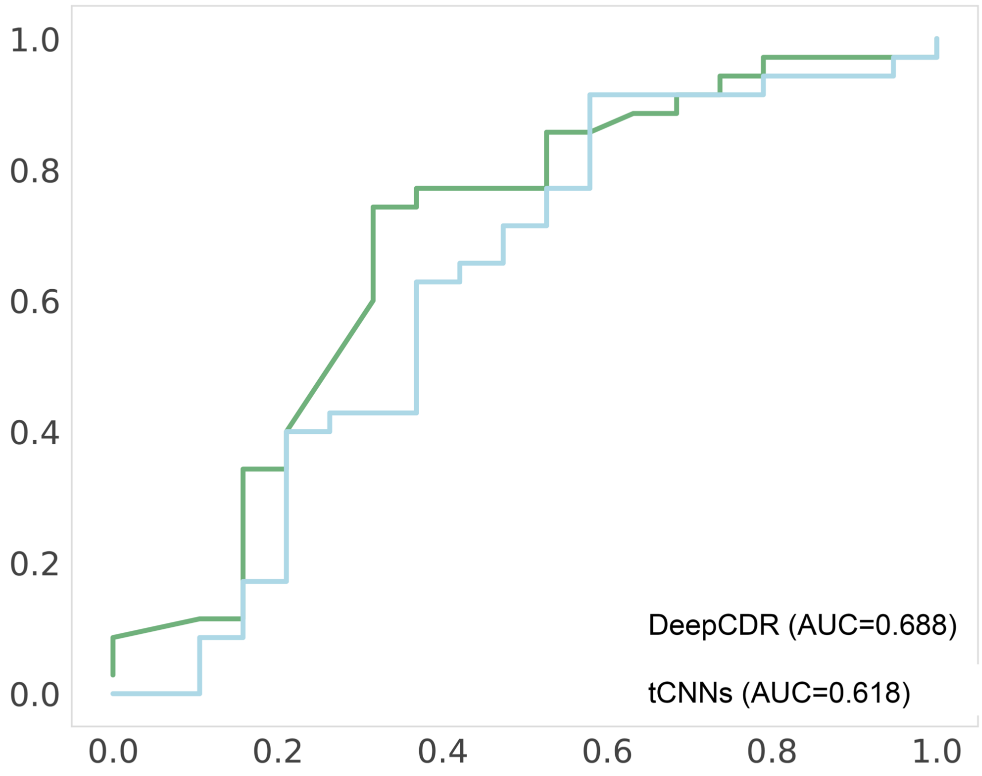


**Supplementary Figure 5.** The performance of DeepCDR and tCNNs in external patient data. DeepCDR achieves an AUC of 0.688, compared to 0.618 of tCNNs.

### Supplementary Tables

**Supplementary Table 1**. The detailed hyperparameters of uniform graph convolutional network (UGCN) architecture of DeepCDR model.

| Layers | Output shape | Operation |
|---|---|---|
| Drug_feat_input | $\mathrm{None} \times 100 \times 75$ | |
| Drug_adj_input | $\mathrm{None} \times 100 \times 100$ | |
| Graph_conv1 | $\mathrm{None} \times 100 \times 256$ | Kernel size = (75,256) |
| | | ReLu activation |
| | | Batch normalization |
| | | Dropout rate=0.1 |
| Graph_conv2 | $\mathrm{None} \times 100 \times 256$ | Kernel size = (256,256) |
| | | ReLu activation |
| | | Batch normalization |
| | | Dropout rate=0.1 |
| Graph_conv3 | $\mathrm{None} \times 100 \times 100$ | Kernel size = (256,100) |
| | | ReLu activation |
| | | Batch normalization |
| | | Dropout rate=0.1 |
| Global max pooling | $\mathrm{None} \times 100$ | |

**Supplementary Table 2**. The detailed hyperparameters of three subnetworks from DeepCDR model. These three subnetworks aim at extracting high-level features of genomic, transcriptomic and epigenomic profile of cancer cell line, respectively.

| Layers | Output shape | Operation |
|---|---|---|
| Genomic input | $\mathrm{None} \times 34673$ | |
| Reshape layer | $\mathrm{None} \times 1 \times 34673 \times 1$ | |
| Convolution | $\mathrm{None} \times 1 \times 6795 \times 50$ | $1 \times 700$ conv, stride=5 |
| | | Tanh activation |
| Max pooling | $\mathrm{None} \times 1 \times 1359 \times 50$ | Pooling size = (1,5) |
| Convolution | $\mathrm{None} \times 1 \times 678 \times 30$ | $1 \times 5$ conv, stride=2 |
| | | Relu activation |
| Max pooling | $\mathrm{None} \times 1 \times 67 \times 30$ | Pooling size = (1,10) |
| Flatten | $\mathrm{None} \times 2010$ | |
| Fully connected layer | $\mathrm{None} \times 100$ | Relu activation, dropout rate=0.1 |
| Transcriptomic input | $\mathrm{None} \times 697$ | |
| Fully connected layer | $\mathrm{None} \times 256$ | Tanh activation, batch normalization, dropout rate=0.1 |
| Fully connected layer | $\mathrm{None} \times 100$ | ReLu activation |
| Epigenomic input | $\mathrm{None} \times 808$ | |
| Fully connected layer | $\mathrm{None} \times 256$ | Tanh activation, batch normalization, dropout rate=0.1 |
| Fully connected layer | $\mathrm{None} \times 100$ | ReLu activation |
| Concatenation | $\mathrm{None} \times 300$ | |

**Supplementary Table 3**. The 238 drugs used in our study after filtering the drugs without PubChem id. Note that several drugs with different GDSC ids may share the same PubChem ids due to the different screening condition. We treated them as individual drugs in our study.

| GDSC id | PubChem id | Drug name | GDSC id | PubChem id | Drug name |
|---|---|---|---|---|---|
| 1 | 176870 | Erlotinib | 282 | 6445562 | Pelitinib |
| 3 | 5384616 | Rapamycin | 283 | 25167777 | Omipalisib |
| 5 | 5329102 | Sunitinib | 287 | 56965967 | KIN001-244 |
| 6 | 10461815 | PHA-665752 | 288 | 3796 | WHI-P97 |
| 9 | 462382 | MG-132 | 290 | 10451420 | KIN001-260 |
| 11 | 36314 | Paclitaxel | 291 | 44143370 | KIN001-266 |
| 29 | 11676786 | AZ628 | 292 | 10074640 | Masitinib |
| 30 | 216239 | Sorafenib | 293 | 11282283 | Amuvatinib |
| 32 | 5494449 | Tozasertib | 294 | 25195352 | MPS-1-IN-1 |
| 34 | 5291 | Imatinib | 295 | 16747388 | NVP-BHG712 |
| 35 | 16038120 | NVP-TAE684 | 298 | 9868037 | OSI-930 |
| 37 | 11626560 | Crizotinib | 299 | 44224160 | OSI-027 |
| 38 | 10302451 | Saracatinib | 300 | 25257557 | CX-5461 |
| 41 | 76044 | S-Trityl-L-cysteine | 301 | 46191454 | PHA-793887 |
| 45 | 16760646 | Z-LLNle-CHO | 302 | 9884685 | PI-103 |
| 51 | 3062316 | Dasatinib | 303 | 6852167 | PIK-93 |
| 52 | 5311510 | GNF-2 | 304 | 9967941 | SB52334 |
| 53 | 644215 | CGP-60474 | 305 | 9903786 | TPCA-1 |
| 54 | 24825971 | CGP-082996 | 306 | 16722836 | Fedratinib |
| 55 | 9549184 | A-770041 | 308 | 42642645 | Foretinib |
| 56 | 11844351 | WH-4-023 | 309 | 9810884 | Y-39983 |
| 59 | 49821040 | WZ-1-84 | 310 | 9956222 | YM201636 |
| 60 | 11364421 | BI-2536 | 312 | 9911830 | Tivozanib |
| 62 | 10390396 | BMS-536924 | 326 | 16725726 | GSK690693 |
| 63 | 20635522 | BMS-509744 | 328 | 24772860 | SNX-2112 |
| 64 | 16663089 | CMK | 333 | 447912 | T0901317 |
| 71 | 4993 | Pyrimethamine | 341 | 5113032 | Selisistat |
| 83 | 49836027 | JW-7-52-1 | 345 | 66577006 | KIN001-270 |
| 86 | 10172943 | A-443654 | 1001 | 65110 | AICA Ribonucleotide |
| 87 | 9826308 | GW843682X | 1004 | 6710780 | Vinblastine |
| 88 | 4261 | Entinostat | 1005 | 84691 | Cisplatin |
| 89 | 7251185 | Parthenolide | 1006 | 6253 | Cytarabine |
| 91 | 11626927 | GSK319347A | 1007 | 148124 | Docetaxel |
| 94 | 9907093 | TGX221 | 1008 | 126941 | Methotrexate |
| 104 | 387447 | Bortezomib | 1009 | 444795 | Tretinoin |
| 106 | 46844147 | XMD8-85 | 1010 | 123631 | Gefitinib |
| 110 | 160355 | Seliciclib | 1011 | 24978538 | Navitoclax |
| 111 | 5717801 | Salubrinal | 1012 | 5311 | Vorinostat |
| 119 | 208908 | Lapatinib | 1013 | 644241 | Nilotinib |

| | | | | | |
|---|---|---|---|---|---|
| 127 | 16095342 | GSK269962A | 1014 | 44182295 | Refametinib |
| 133 | 31703 | Doxorubicin | 1015 | 6918454 | CI-1040 |
| 134 | 36462 | Etoposide | 1016 | 6918289 | Temsirolimus |
| 135 | 60750 | Gemcitabine | 1017 | 23725625 | Olaparib |
| 136 | 5746 | Mitomycin-C | 1018 | 11960529 | Veliparib |
| 140 | 5311497 | Vinorelbine | 1019 | 5328940 | Bosutinib |
| 147 | 5459322 | NSC-87877 | 1020 | 216326 | Lenalidomide |
| 150 | 2375 | Bicalutamide | 1021 | 6450551 | Axitinib |
| 151 | 4263900 | QS11 | 1022 | 11152667 | AZD7762 |
| 152 | 44551660 | CP466722 | 1023 | 9943465 | GW441756 |
| 154 | 9956119 | CHIR-99021 | 1024 | 126565 | Lestaurtinib |
| 155 | 24826799 | Ponatinib | 1025 | 176158 | SB216763 |
| 156 | 44137675 | AZD6482 | 1026 | 6505803 | Tanespimycin |
| 157 | 25222038 | JNK-9L | 1028 | 10341154 | VX-702 |
| 158 | 11713159 | PF-562271 | 1029 | 11667893 | Motesanib |
| 159 | 53302361 | HG6-64-1 | 1030 | 5278396 | KU-55933 |
| 163 | 46907787 | JQ1 | 1031 | 300471 | Elesclomol |
| 165 | 560326 | DMOG | 1032 | 10184653 | Afatinib |
| 166 | 3005532 | FTI-277 | 1033 | 24776445 | Vismodegib |
| 167 | 10027278 | OSU-03012 | 1036 | 24180719 | PLX-4720 |
| 170 | 5208 | Shikonin | 1037 | 10077147 | BX795 |
| 171 | 10196499 | AKT inhibitor VIII | 1038 | 11327430 | NU7441 |
| 172 | 3218 | Embelin | 1039 | 10459196 | SL0101 |
| 173 | 3463933 | FH535 | 1042 | 156422 | Doramapimod |
| 175 | 6753378 | PAC-1 | 1043 | 11624601 | JNK Inhibitor VIII |
| 176 | 521106 | IPA-3 | 1046 | 10384072 | Wee1 Inhibitor |
| 177 | 25022668 | GSK650394 | 1047 | 11433190 | Nutlin-3a (-) |
| 178 | 10200390 | BAY-61-3606 | 1049 | 1401 | PD173074 |
| 179 | 3385 | 5-Fluorouracil | 1050 | 9914412 | ZM447439 |
| 180 | 446378 | Thapsigargin | 1052 | 44450571 | RO-3306 |
| 182 | 11404337 | Obatoclax Mesylate | 1053 | 46930998 | MK-2206 |
| 184 | 24785538 | BMS-754807 | 1054 | 5330286 | Palbociclib |
| 185 | 11640390 | Linsitinib | 1057 | 11977753 | Dactolisib |
| 186 | 82146 | Bexarotene | 1058 | 17755052 | Pictilisib |
| 192 | 54676905 | LFM-A13 | 1059 | 25262965 | AZD8055 |
| 193 | 11617559 | GW-2580 | 1060 | 9826528 | PD0325901 |
| 194 | 10096043 | Luminespib | 1061 | 11316960 | SB590885 |
| 196 | 8249 | Phenformin | 1062 | 10127622 | Selumetinib |
| 197 | 5280757 | Bryostatin 1 | 1066 | 44137675 | AZD6482 |
| 199 | 10113978 | Pazopanib | 1067 | 2314623 | CCT007093 |
| 200 | 6445533 | Dacinostat | 1069 | 9938202 | EHT-1864 |
| 201 | 448013 | Epothilone B | 1072 | 46883536 | Avagacestat |
| 202 | 25124816 | GSK1904529A | 1091 | 10390396 | BMS-536924 |

| | | | | | |
|---|---|---|---|---|---|
| 203 | 9813758 | BMS-345541 | 1114 | 85668777 | Cetuximab |
| 204 | 159324 | Tipifarnib | 1129 | 51371303 | PF-4708671 |
| 205 | 46883536 | Avagacestat | 1133 | 11609586 | Serdemetan |
| 206 | 25126798 | Ruxolitinib | 1149 | 11455910 | TW 37 |
| 207 | 10109823 | AS601245 | 1164 | 46843772 | XMD8-92 |
| 208 | 6450816 | Ispinesib Mesylate | 1170 | 5327091 | CCT-018159 |
| 219 | 11338033 | AT-7519 | 1175 | 9931953 | Rucaparib |
| 221 | 9952773 | TAK-715 | 1192 | 16095342 | GSK269962A |
| 222 | 11754511 | BX-912 | 1194 | 9858940 | SB505124 |
| 223 | 11647372 | ZSTK474 | 1199 | 2733526 | Tamoxifen |
| 224 | 5289247 | AS605240 | 1218 | 46907787 | JQ1 |
| 226 | 46885626 | GSK1070916 | 1219 | 71271629 | PFI-1 |
| 228 | 10196499 | AKT inhibitor VIII | 1230 | 54685215 | IOX2 |
| 229 | 176167 | Enzastaurin | 1236 | 46224516 | UNC0638 |
| 230 | 11373846 | GSK429286A | 1239 | 44632017 | YK-4-279 |
| 238 | 11625818 | Idelalisib | 1241 | 9956119 | CHIR-99021 |
| 245 | 46224516 | UNC0638 | 1242 | 9863776 | (5Z)-7-Oxozeaenol |
| 249 | 25102847 | Cabozantinib | 1243 | 637858 | Piperlongumine |
| 254 | 24889392 | Quizartinib | 1248 | 6914657 | Daporinad |
| 255 | 9874913 | CP724714 | 1259 | 44819241 | Talazoparib |
| 258 | 704473 | STF-62247 | 1262 | 57339144 | UNC1215 |
| 260 | 53340664 | NG-25 | 1264 | 56962337 | SGC0946 |
| 262 | 11634725 | VX-11e | 1268 | 2726824 | XAV939 |
| 263 | 11493598 | FR-180204 | 1371 | 24180719 | PLX-4720 |
| 265 | 53394750 | Tubastatin A | 1372 | 11707110 | Trametinib |
| 266 | 9910224 | Zibotentan | 1373 | 44462760 | Dabrafenib |
| 268 | 11178236 | Sepantronium bromide | 1375 | 5394 | Temozolomide |
| 269 | 42640 | NSC-207895 | 1377 | 10184653 | Afatinib |
| 271 | 24894414 | VNLG/124 | 1378 | 5460769 | Bleomycin |
| 272 | 6918848 | AR-42 | 1494 | 104842 | SN-38 |
| 273 | 24756910 | CUDC-101 | 1495 | 23725625 | Olaparib |
| 274 | 6918638 | Belinostat | 1498 | 10127622 | Selumetinib |
| 275 | 46943432 | I-BET-762 | 1502 | 2375 | Bicalutamide |
| 276 | 24951314 | CAY10603 | 1526 | 44182295 | Refametinib |
| 277 | 11485656 | Linifanib | 1527 | 17755052 | Pictilisib |
| 279 | 46931012 | BIX02189 | 1529 | 16720766 | Pevonedistat |
| 281 | 49806720 | Alectinib | 1530 | 78243717 | PFI-3 |

**Supplementary Table 4**. The 561 cancer cell lines that were used in our study.

| Demap id | Cell line name | TCGA cancer type | Demap id | Cell line name | TCGA cancer type |
|---|---|---|---|---|---|
| ACH-000001 | NIH:OVCAR-3 | OV | ACH-000532 | SNU-61 | COAD/READ |
| ACH-000002 | HL-60 | LAML | ACH-000534 | WSU-DLCL2 | DLBC |
| ACH-000004 | HEL | LAML | ACH-000535 | BxPC-3 | PAAD |
| ACH-000006 | MONO-MAC-6 | LAML | ACH-000536 | BT-20 | BRCA |
| ACH-000007 | LS513 | COAD/READ | ACH-000544 | OE21 | ESCA |
| ACH-000008 | A101D | SKCM | ACH-000545 | VM-CUB1 | BLCA |
| ACH-000009 | C2BBe1 | COAD/READ | ACH-000546 | HSC-4 | HNSC |
| ACH-000012 | HCC827 | LUAD | ACH-000547 | HT-1197 | BLCA |
| ACH-000018 | T24 | BLCA | ACH-000548 | BHY | HNSC |
| ACH-000019 | MCF7 | BRCA | ACH-000551 | K-562 | LCML |
| ACH-000020 | MHH-CALL-2 | ALL | ACH-000554 | UACC-893 | BRCA |
| ACH-000023 | PA-TU-8988T | PAAD | ACH-000555 | A-498 | KIRC |
| ACH-000024 | OPM-2 | MM | ACH-000558 | A172 | GBM |
| ACH-000035 | NCI-H1650 | LUAD | ACH-000559 | NCI-H1836 | SCLC |
| ACH-000038 | EHEB | CLL | ACH-000560 | ECC10 | STAD |
| ACH-000040 | U-118 MG | GBM | ACH-000561 | NA | ESCA |
| ACH-000042 | Panc 02.03 | PAAD | ACH-000562 | HCC-78 | LUAD |
| ACH-000046 | ACHN | KIRC | ACH-000563 | EBC-1 | LUSC |
| ACH-000047 | GCIY | STAD | ACH-000565 | RCM-1 | COAD/READ |
| ACH-000048 | TOV-112D | OV | ACH-000566 | SW-1710 | BLCA |
| ACH-000050 | NCI-H929 | MM | ACH-000567 | ST486 | DLBC |
| ACH-000052 | A-673 | SARC | ACH-000568 | UACC-812 | BRCA |
| ACH-000054 | HT-1080 | SARC | ACH-000570 | YKG1 | GBM |
| ACH-000055 | D283 Med | MB | ACH-000572 | G-361 | SKCM |
| ACH-000056 | DOHH-2 | DLBC | ACH-000573 | MDA-MB-436 | BRCA |
| ACH-000058 | ML-1 | THCA | ACH-000574 | FU-OV-1 | OV |
| ACH-000059 | SUP-B15 | ALL | ACH-000579 | UACC-257 | SKCM |
| ACH-000060 | Panc 10.05 | PAAD | ACH-000580 | C32 | SKCM |
| ACH-000061 | HH | ALL | ACH-000585 | EPLC-272H | LUSC |
| ACH-000062 | RERF-LC-MS | LUAD | ACH-000586 | NCI-H1876 | SCLC |
| ACH-000065 | OCI-AML5 | LAML | ACH-000587 | NCI-H1975 | LUAD |
| ACH-000067 | Hs 683 | LGG | ACH-000589 | NCI-H1437 | LUAD |
| ACH-000070 | 697 | ALL | ACH-000595 | LN-229 | GBM |
| ACH-000072 | MEG-01 | LCML | ACH-000596 | LCLC-97TM1 | NA |
| ACH-000073 | GRANTA-519 | DLBC | ACH-000599 | PA-TU-8902 | PAAD |
| ACH-000074 | KU812 | LCML | ACH-000603 | BEN | UNABLE TO CLASSIFY |
| ACH-000075 | U-87 MG | GBM | ACH-000605 | TE-6 | ESCA |
| ACH-000078 | MHH-NB-11 | NB | ACH-000607 | KYM-1 | SARC |
| ACH-000081 | GDM-1 | LAML | ACH-000609 | SF126 | GBM |
| ACH-000082 | G-292, clone A141B1 | SARC | ACH-000610 | NCI-H2227 | SCLC |

| | | | | | |
|---|---|---|---|---|---|
| ACH-000087 | SK-ES-1 | SARC | ACH-000611 | SU-DHL-6 | DLBC |
| ACH-000090 | PC-3 | PRAD | ACH-000613 | HOS | SARC |
| ACH-000091 | OV56 | OV | ACH-000614 | RVH-421 | SKCM |
| ACH-000092 | NCI-H2452 | MESO | ACH-000615 | SK-MEL-28 | SKCM |
| ACH-000094 | HPAF-II | PAAD | ACH-000616 | Hs 746T | STAD |
| ACH-000096 | G-401 | SARC | ACH-000617 | OVCAR-4 | OV |
| ACH-000098 | GAMG | LGG | ACH-000619 | PE/CA-PJ15 | HNSC |
| ACH-000099 | SIMA | NB | ACH-000621 | MDA-MB-157 | BRCA |
| ACH-000102 | GMS-10 | GBM | ACH-000624 | HCC1806 | BRCA |
| ACH-000103 | Caov-4 | OV | ACH-000625 | Hep 3B2.1-7 | LIHC |
| ACH-000104 | Loucy | ALL | ACH-000626 | U266B1 | MM |
| ACH-000105 | ALL-SIL | ALL | ACH-000627 | LCLC-103H | NA |
| ACH-000106 | JVM-2 | CLL | ACH-000628 | NCI-H596 | LUAD |
| ACH-000107 | Capan-2 | PAAD | ACH-000633 | FU97 | STAD |
| ACH-000111 | HCC1187 | BRCA | ACH-000636 | RPMI-8402 | ALL |
| ACH-000112 | SIG-M5 | LAML | ACH-000637 | KYSE-520 | ESCA |
| ACH-000113 | OCI-AML2 | LAML | ACH-000638 | NCI-H441 | LUAD |
| ACH-000114 | SU.86.86 | PAAD | ACH-000639 | NCI-H211 | SCLC |
| ACH-000115 | VCaP | PRAD | ACH-000640 | SK-MEL-31 | SKCM |
| ACH-000116 | OAW28 | OV | ACH-000641 | CMK | LAML |
| ACH-000117 | EFM-192A | BRCA | ACH-000643 | HDQ-P1 | BRCA |
| ACH-000118 | HUP-T3 | PAAD | ACH-000644 | COLO 829 | SKCM |
| ACH-000120 | CHP-212 | NB | ACH-000647 | TE-1 | ESCA |
| ACH-000121 | NCI-H2405 | LUAD | ACH-000648 | NCI-H28 | MESO |
| ACH-000124 | OCI-LY-19 | DLBC | ACH-000649 | 786-O | KIRC |
| ACH-000129 | NCI-H1341 | SCLC | ACH-000650 | IGR-37 | SKCM |
| ACH-000132 | JHOS-2 | OV | ACH-000651 | SW620 | COAD/READ |
| ACH-000135 | Hs 940.T | SKCM | ACH-000652 | SUIT-2 | PAAD |
| ACH-000136 | CHP-126 | NB | ACH-000654 | Raji | DLBC |
| ACH-000137 | 8-MG-BA | GBM | ACH-000655 | SF268 | GBM |
| ACH-000138 | CFPAC-1 | PAAD | ACH-000656 | SU-DHL-8 | DLBC |
| ACH-000139 | Panc 03.27 | PAAD | ACH-000657 | A2780 | UNABLE TO CLASSIFY |
| ACH-000142 | CAL-29 | BLCA | ACH-000660 | SU-DHL-5 | DLBC |
| ACH-000144 | RERF-GC-1B | STAD | ACH-000662 | COR-L23 | NA |
| ACH-000146 | THP-1 | LAML | ACH-000663 | OVTOKO | OV |
| ACH-000147 | T-47D | BRCA | ACH-000664 | SU-DHL-1 | NA |
| ACH-000148 | Hs 578T | BRCA | ACH-000665 | SK-MES-1 | LUSC |
| ACH-000149 | SK-N-SH | NB | ACH-000667 | HCC-44 | LUAD |
| ACH-000151 | JM1 | DLBC | ACH-000668 | HCC70 | BRCA |
| ACH-000153 | NCI-H2052 | MESO | ACH-000669 | SW 900 | LUSC |
| ACH-000155 | SW 1990 | PAAD | ACH-000670 | SBC-5 | SCLC |
| ACH-000157 | A4/Fuk | DLBC | ACH-000672 | IA-LM | NA |
| ACH-000159 | OS-RC-2 | KIRC | ACH-000674 | NUGC-4 | STAD |

| | | | | | |
|---|---|---|---|---|---|
| ACH-000161 | COR-L105 | LUAD | ACH-000677 | SW 1573 | LUAD |
| ACH-000162 | GA-10 | DLBC | ACH-000678 | MKN7 | STAD |
| ACH-000168 | NOMO-1 | LAML | ACH-000679 | OE19 | ESCA |
| ACH-000169 | RD | SARC | ACH-000680 | SW948 | COAD/READ |
| ACH-000171 | VMRC-RCZ | KIRC | ACH-000681 | A549 | LUAD |
| ACH-000174 | CAL-62 | THCA | ACH-000684 | KMRC-1 | KIRC |
| ACH-000176 | LOU-NH91 | LUSC | ACH-000688 | OV7 | OV |
| ACH-000178 | Hs 766T | PAAD | ACH-000689 | RH-18 | SARC |
| ACH-000181 | SCC-9 | HNSC | ACH-000691 | HCC2157 | BRCA |
| ACH-000183 | L-363 | MM | ACH-000693 | KYSE-180 | ESCA |
| ACH-000186 | NCI-H2444 | UNABLE TO CLASSIFY | ACH-000694 | TE-9 | ESCA |
| ACH-000187 | COR-L311 | SCLC | ACH-000696 | OVCAR-8 | OV |
| ACH-000188 | SCC-25 | HNSC | ACH-000697 | A3/KAW | DLBC |
| ACH-000189 | RCC10RGB | KIRC | ACH-000698 | DMS 53 | SCLC |
| ACH-000190 | HD-MY-Z | NA | ACH-000699 | HCC1395 | BRCA |
| ACH-000191 | BHT-101 | THCA | ACH-000702 | L-1236 | NA |
| ACH-000196 | HCC1599 | BRCA | ACH-000703 | DMS 79 | SCLC |
| ACH-000197 | TALL-1 | ALL | ACH-000704 | OAW42 | OV |
| ACH-000198 | EOL-1 | LAML | ACH-000706 | EKVX | LUAD |
| ACH-000200 | NMC-G1 | LGG | ACH-000713 | Caov-3 | OV |
| ACH-000201 | A-204 | SARC | ACH-000714 | KMS-11 | MM |
| ACH-000204 | LP-1 | MM | ACH-000716 | TT2609-C02 | THCA |
| ACH-000207 | Detroit 562 | HNSC | ACH-000717 | COLO-680N | ESCA |
| ACH-000210 | CADO-ES1 | SARC | ACH-000718 | NCI-H2291 | LUAD |
| ACH-000211 | Daoy | MB | ACH-000719 | RMG-I | OV |
| ACH-000212 | CAL-120 | BRCA | ACH-000720 | TCCSUP | BLCA |
| ACH-000218 | PL-21 | LAML | ACH-000722 | SNU-C1 | COAD/READ |
| ACH-000219 | A-375 | SKCM | ACH-000725 | HCC202 | BRCA |
| ACH-000221 | SNU-398 | LIHC | ACH-000727 | NCI-H2066 | SCLC |
| ACH-000222 | AsPC-1 | PAAD | ACH-000729 | NCI-H1963 | SCLC |
| ACH-000223 | HCC1937 | BRCA | ACH-000730 | SK-MEL-5 | SKCM |
| ACH-000225 | ECC12 | STAD | ACH-000733 | NCI-H1838 | LUAD |
| ACH-000226 | SUP-M2 | NA | ACH-000738 | GB-1 | GBM |
| ACH-000227 | KP-N-YN | NB | ACH-000740 | A-253 | HNSC |
| ACH-000228 | BICR 31 | HNSC | ACH-000743 | COR-L95 | SCLC |
| ACH-000231 | KALS-1 | LGG | ACH-000744 | NCI-H1623 | LUAD |
| ACH-000232 | U-251 MG | GBM | ACH-000745 | MOLP-8 | MM |
| ACH-000235 | Panc 04.03 | PAAD | ACH-000748 | SJSA-1 | SARC |
| ACH-000236 | SW1417 | COAD/READ | ACH-000749 | DMS 273 | SCLC |
| ACH-000238 | SCC-4 | HNSC | ACH-000750 | LOX IMVI | SKCM |
| ACH-000242 | RT4 | BLCA | ACH-000751 | OCI-M1 | LAML |
| ACH-000243 | DAN-G | PAAD | ACH-000752 | NCI-H196 | SCLC |
| ACH-000244 | DK-MG | GBM | ACH-000754 | L-428 | NA |

| | | | | | |
|---|---|---|---|---|---|
| ACH-000245 | BL-41 | DLBC | ACH-000755 | HCC2218 | BRCA |
| ACH-000247 | OCUM-1 | STAD | ACH-000756 | GI-1 | LGG |
| ACH-000248 | AU565 | BRCA | ACH-000759 | MDA-MB-175-VII | BRCA |
| ACH-000249 | CL-11 | COAD/READ | ACH-000763 | MM1-S | MM |
| ACH-000250 | KMRC-20 | KIRC | ACH-000766 | NCI-H1648 | LUAD |
| ACH-000254 | SCC-15 | HNSC | ACH-000767 | NCI-H526 | SCLC |
| ACH-000257 | COR-L279 | SCLC | ACH-000768 | MDA-MB-231 | BRCA |
| ACH-000258 | DU4475 | BRCA | ACH-000770 | P31/FUJ | LAML |
| ACH-000260 | SK-N-AS | NB | ACH-000772 | TE 441.T | SARC |
| ACH-000264 | Calu-6 | LUAD | ACH-000775 | NCI-H727 | NA |
| ACH-000267 | HDLM-2 | NA | ACH-000776 | ONS-76 | MB |
| ACH-000269 | AM-38 | GBM | ACH-000778 | HSC-3 | HNSC |
| ACH-000270 | HPAC | PAAD | ACH-000780 | NCI-H1105 | SCLC |
| ACH-000271 | SU-DHL-10 | DLBC | ACH-000781 | NCI-H2023 | LUAD |
| ACH-000273 | SF539 | LGG | ACH-000783 | CAMA-1 | BRCA |
| ACH-000277 | HCC1419 | BRCA | ACH-000784 | KYSE-70 | ESCA |
| ACH-000287 | NU-DUL-1 | DLBC | ACH-000786 | Daudi | DLBC |
| ACH-000288 | BT-549 | BRCA | ACH-000787 | LXF-289 | LUAD |
| ACH-000290 | NCI-H209 | SCLC | ACH-000788 | A2058 | SKCM |
| ACH-000291 | OV-90 | OV | ACH-000789 | NCI-H810 | NA |
| ACH-000292 | NCI-H841 | SCLC | ACH-000790 | SHP-77 | SCLC |
| ACH-000294 | NB-4 | LAML | ACH-000792 | BFTC-909 | KIRC |
| ACH-000295 | EM-2 | LCML | ACH-000793 | KATO III | STAD |
| ACH-000298 | NCI-H2029 | SCLC | ACH-000794 | BICR 22 | HNSC |
| ACH-000301 | LAMA-84 | LCML | ACH-000795 | MOLT-13 | ALL |
| ACH-000303 | SNU-5 | STAD | ACH-000798 | CL-40 | COAD/READ |
| ACH-000304 | WM-115 | SKCM | ACH-000800 | NCI-H446 | SCLC |
| ACH-000305 | EC-GI-10 | ESCA | ACH-000802 | BFTC-905 | BLCA |
| ACH-000308 | EFO-21 | OV | ACH-000803 | COLO 668 | SCLC |
| ACH-000309 | SK-LU-1 | LUAD | ACH-000804 | NB-1 | NB |
| ACH-000311 | NCI-H2122 | LUAD | ACH-000805 | COLO-679 | SKCM |
| ACH-000315 | KARPAS-422 | DLBC | ACH-000806 | L-540 | NA |
| ACH-000318 | TE-10 | ESCA | ACH-000810 | SK-MEL-30 | SKCM |
| ACH-000320 | PSN1 | PAAD | ACH-000811 | SK-OV-3 | OV |
| ACH-000322 | HT-144 | SKCM | ACH-000812 | COLO-783 | SKCM |
| ACH-000323 | 42-MG-BA | GBM | ACH-000814 | Hs 939.T | SKCM |
| ACH-000326 | JURL-MK1 | LCML | ACH-000815 | KM-H2 | NA |
| ACH-000327 | NCI-H1395 | LUAD | ACH-000816 | NCI-H524 | SCLC |
| ACH-000329 | CCF-STTG1 | GBM | ACH-000817 | RPMI 8226 | MM |
| ACH-000330 | EFM-19 | BRCA | ACH-000818 | BT-483 | BRCA |
| ACH-000332 | YAPC | PAAD | ACH-000821 | EJM | MM |
| ACH-000334 | DB | DLBC | ACH-000822 | SK-MEL-24 | SKCM |
| ACH-000335 | MSTO-211H | MESO | ACH-000823 | KYSE-140 | ESCA |

| | | | | | |
|---|---|---|---|---|---|
| ACH-000336 | OCI-AML3 | LAML | ACH-000824 | KYSE-510 | ESCA |
| ACH-000341 | SK-N-FI | NB | ACH-000825 | HOP-92 | NA |
| ACH-000343 | NCI-H522 | LUAD | ACH-000826 | CAL-12T | UNABLE TO CLASSIFY |
| ACH-000345 | KP-N-RT-BM-1 | NB | ACH-000827 | WM-793 | SKCM |
| ACH-000346 | JVM-3 | CLL | ACH-000828 | ZR-75-30 | BRCA |
| ACH-000347 | QGP-1 | PAAD | ACH-000830 | NCI-H1436 | SCLC |
| ACH-000348 | RPMI-7951 | SKCM | ACH-000832 | CAL 27 | HNSC |
| ACH-000349 | HCC1500 | BRCA | ACH-000835 | GCT | SARC |
| ACH-000350 | COLO-678 | COAD/READ | ACH-000838 | AMO-1 | MM |
| ACH-000351 | MKN1 | STAD | ACH-000841 | NCI-H2087 | LUAD |
| ACH-000352 | HCC1428 | BRCA | ACH-000843 | HARA | LUSC |
| ACH-000353 | TE-15 | ESCA | ACH-000846 | FaDu | HNSC |
| ACH-000355 | NCI-H82 | SCLC | ACH-000847 | HGC-27 | STAD |
| ACH-000356 | MKN-45 | STAD | ACH-000849 | MDA-MB-468 | BRCA |
| ACH-000357 | JeKo-1 | DLBC | ACH-000852 | NCI-H1435 | LUAD |
| ACH-000358 | NCI-H69 | SCLC | ACH-000853 | NCI-H661 | NA |
| ACH-000359 | MG-63 | SARC | ACH-000855 | KYSE-150 | ESCA |
| ACH-000360 | NCI-H508 | COAD/READ | ACH-000856 | CAL-51 | BRCA |
| ACH-000361 | SK-HEP-1 | LIHC | ACH-000857 | CAL-85-1 | BRCA |
| ACH-000362 | MOLM-13 | LAML | ACH-000858 | KNS-62 | LUSC |
| ACH-000363 | SK-MM-2 | MM | ACH-000859 | HCC1954 | BRCA |
| ACH-000364 | U-2 OS | SARC | ACH-000860 | NCI-H358 | LUAD |
| ACH-000365 | SU-DHL-4 | DLBC | ACH-000861 | HOP-62 | LUAD |
| ACH-000366 | SK-N-DZ | NB | ACH-000863 | DBTRG-05MG | GBM |
| ACH-000369 | MOLM-16 | LAML | ACH-000864 | COLO 684 | UCEC |
| ACH-000371 | RL | DLBC | ACH-000866 | NCI-H1048 | SCLC |
| ACH-000372 | P12-ICHIKAWA | ALL | ACH-000867 | ChaGo-K-1 | UNABLE TO CLASSIFY |
| ACH-000373 | SKM-1 | LAML | ACH-000869 | NCI-H1568 | LUAD |
| ACH-000374 | HCC1143 | BRCA | ACH-000871 | NCI-H510 | SCLC |
| ACH-000375 | G-402 | SARC | ACH-000874 | RS4;11 | ALL |
| ACH-000376 | SF-295 | GBM | ACH-000875 | NCI-H2347 | LUAD |
| ACH-000378 | NCI-H647 | LUAD | ACH-000876 | MDA-MB-415 | BRCA |
| ACH-000379 | NCI-H1781 | LUAD | ACH-000878 | HCC-15 | LUSC |
| ACH-000380 | KMS-12-BM | MM | ACH-000879 | MFE-296 | UCEC |
| ACH-000381 | T84 | COAD/READ | ACH-000880 | AGS | STAD |
| ACH-000383 | OE33 | ESCA | ACH-000881 | MEL-JUSO | SKCM |
| ACH-000384 | SW 780 | BLCA | ACH-000882 | IGR-1 | SKCM |
| ACH-000386 | KG-1 | LAML | ACH-000885 | TOV-21G | OV |
| ACH-000389 | H4 | LGG | ACH-000886 | NCI-H2009 | LUAD |
| ACH-000392 | Calu-3 | LUAD | ACH-000888 | NCI-H1793 | LUAD |
| ACH-000394 | NCI-H2081 | SCLC | ACH-000890 | SW 1271 | SCLC |
| ACH-000396 | J82 | BLCA | ACH-000893 | NCI-H1651 | LUAD |
| ACH-000399 | NCI-H2196 | SCLC | ACH-000895 | CL-34 | COAD/READ |

| | | | | | |
|---|---|---|---|---|---|
| ACH-000400 | SK-CO-1 | COAD/READ | ACH-000896 | 647-V | BLCA |
| ACH-000401 | COLO-800 | SKCM | ACH-000900 | NCI-H23 | LUAD |
| ACH-000403 | NCI-H747 | COAD/READ | ACH-000903 | FTC-133 | THCA |
| ACH-000408 | TE-5 | ESCA | ACH-000905 | 5637 | BLCA |
| ACH-000410 | Saos-2 | SARC | ACH-000906 | ES-2 | OV |
| ACH-000411 | 769-P | KIRC | ACH-000910 | MDA-MB-453 | BRCA |
| ACH-000414 | NCI-H1944 | LUAD | ACH-000911 | NUGC-3 | STAD |
| ACH-000416 | NCI-H838 | LUAD | ACH-000913 | ESS-1 | UCEC |
| ACH-000417 | Panc 08.13 | PAAD | ACH-000914 | HT | DLBC |
| ACH-000420 | SNU-449 | LIHC | ACH-000915 | IPC-298 | SKCM |
| ACH-000421 | SW837 | COAD/READ | ACH-000917 | TE-4 | ESCA |
| ACH-000422 | SNU-475 | LIHC | ACH-000918 | MOLT-16 | ALL |
| ACH-000423 | SK-MEL-3 | SKCM | ACH-000920 | CML-T1 | LCML |
| ACH-000425 | UACC-62 | SKCM | ACH-000922 | RCH-ACV | ALL |
| ACH-000427 | NCI-N87 | STAD | ACH-000924 | NCI-H2172 | UNABLE TO CLASSIFY |
| ACH-000429 | A-704 | KIRC | ACH-000927 | BT-474 | BRCA |
| ACH-000430 | TYK-nu | UNABLE TO CLASSIFY | ACH-000929 | NCI-H2110 | UNABLE TO CLASSIFY |
| ACH-000432 | BV-173 | LCML | ACH-000930 | HCC1569 | BRCA |
| ACH-000433 | Caki-1 | KIRC | ACH-000932 | SNU-1 | STAD |
| ACH-000434 | NCI-H1915 | NA | ACH-000934 | MDA-MB-361 | BRCA |
| ACH-000437 | SW 1088 | GBM | ACH-000935 | MDST8 | COAD/READ |
| ACH-000438 | LU65 | NA | ACH-000936 | EFO-27 | OV |
| ACH-000439 | ME-1 | LAML | ACH-000937 | PF-382 | ALL |
| ACH-000440 | CA46 | DLBC | ACH-000938 | NALM-6 | ALL |
| ACH-000441 | SH-4 | SKCM | ACH-000939 | SK-UT-1 | SARC |
| ACH-000443 | OVKATE | OV | ACH-000943 | RKO | COAD/READ |
| ACH-000447 | NCI-H2228 | LUAD | ACH-000944 | NAMALWA | DLBC |
| ACH-000449 | MES-SA | SARC | ACH-000945 | NCI-H650 | LUAD |
| ACH-000450 | MEL-HO | SKCM | ACH-000947 | OVK18 | OV |
| ACH-000451 | NCI-H2085 | LUAD | ACH-000948 | 23132/87 | STAD |
| ACH-000452 | TE-8 | ESCA | ACH-000949 | TGBC11TKB | STAD |
| ACH-000456 | B-CPAP | THCA | ACH-000950 | LoVo | COAD/READ |
| ACH-000457 | CAL-54 | KIRC | ACH-000951 | NCI-H2342 | LUAD |
| ACH-000463 | NCI-H460 | NA | ACH-000953 | SUP-T1 | ALL |
| ACH-000464 | CAS-1 | GBM | ACH-000955 | SNU-407 | COAD/READ |
| ACH-000465 | SK-MEL-1 | SKCM | ACH-000956 | 22Rv1 | PRAD |
| ACH-000467 | HCC-56 | COAD/READ | ACH-000957 | LS 180 | COAD/READ |
| ACH-000469 | YH-13 | GBM | ACH-000958 | SW48 | COAD/READ |
| ACH-000470 | SW1463 | COAD/READ | ACH-000960 | Reh | ALL |
| ACH-000472 | HSC-2 | HNSC | ACH-000963 | CCK-81 | COAD/READ |
| ACH-000473 | RT-112 | BLCA | ACH-000965 | RL95-2 | UCEC |
| ACH-000475 | huH-1 | LIHC | ACH-000966 | IGROV1 | OV |
| ACH-000476 | JHH-4 | LIHC | ACH-000968 | COLO 792 | SKCM |

| | | | | | |
|---|---|---|---|---|---|
| ACH-000478 | SNU-387 | LIHC | ACH-000969 | KM12 | COAD/READ |
| ACH-000480 | HuH-7 | LIHC | ACH-000970 | SNU-C5 | COAD/READ |
| ACH-000481 | NCI-H2170 | LUSC | ACH-000971 | HCT 116 | COAD/READ |
| ACH-000482 | RERF-LC-KJ | LUAD | ACH-000973 | 639-V | BLCA |
| ACH-000483 | SNU-182 | LIHC | ACH-000974 | SNG-M | UCEC |
| ACH-000484 | VMRC-RCW | KIRC | ACH-000976 | HuCCT1 | NA |
| ACH-000486 | KU-19-19 | BLCA | ACH-000978 | EN | UCEC |
| ACH-000488 | TE-11 | ESCA | ACH-000979 | DU 145 | PRAD |
| ACH-000489 | SW1116 | COAD/READ | ACH-000980 | NCI-H1155 | NA |
| ACH-000491 | NCI-H716 | COAD/READ | ACH-000981 | DND-41 | ALL |
| ACH-000493 | SNU-423 | LIHC | ACH-000983 | KCL-22 | LCML |
| ACH-000496 | NCI-H1792 | LUAD | ACH-000985 | LS411N | COAD/READ |
| ACH-000501 | LS123 | COAD/READ | ACH-000986 | HT115 | COAD/READ |
| ACH-000504 | SNB75 | GBM | ACH-000987 | MeWo | SKCM |
| ACH-000505 | RKN | SARC | ACH-000989 | SNU-175 | COAD/READ |
| ACH-000506 | NCI-H146 | SCLC | ACH-000991 | SNU-81 | COAD/READ |
| ACH-000508 | COR-L88 | SCLC | ACH-000995 | JURKAT | ALL |
| ACH-000514 | NCI-H1092 | SCLC | ACH-000997 | HCT-15 | COAD/READ |
| ACH-000515 | HCC-33 | SCLC | ACH-000998 | CW-2 | COAD/READ |
| ACH-000518 | CAL-33 | HNSC | ACH-000999 | SNU-1040 | COAD/READ |
| ACH-000521 | NCI-H2030 | LUAD | ACH-001075 | NCI-H292 | NA |
| ACH-000522 | UM-UC-3 | BLCA | ACH-001106 | KOPN-8 | ALL |
| ACH-000525 | NCI-H2171 | SCLC | ACH-001190 | SK-MEL-2 | SKCM |
| ACH-000527 | OVISE | OV | ACH-001306 | 8305C | THCA |
| ACH-000528 | ABC-1 | LUAD | ACH-001307 | 8505C | THCA |
| ACH-000530 | DMS 114 | SCLC | | | |

**Supplementary Table 5**. The information of TCGA patient data which contains 54 records across 31 patients and 12 drugs. The corresponding clinic responses were also provided. The corresponding gene mutation data, gene expression data and DNA methylation data of the patients were downloaded from Firehose Broad GDAC (http://gdac.broadinstitute.org/runs/stddata__2016_01_28/).

| Patient id | Drug | PubChem ID | DrugBank ID | Measure of response | Label |
|---|---|---|---|---|---|
| TCGA-C5-A1M6 | Bleomycin | 5460769 | DB00290 | Clinical Progressive Disease | 0 |
| TCGA-C5-A1M6 | Vincristine | 13342 | DB00570 | Clinical Progressive Disease | 0 |
| TCGA-C5-A1M6 | Mitomycin | 5746 | DB00305 | Clinical Progressive Disease | 0 |
| TCGA-C5-A1M6 | Hydrocortisone | 5754 | DB00741 | Clinical Progressive Disease | 0 |
| TCGA-FU-A23L | Cisplatin | 84691 | DB00515 | Complete Response | 1 |
| TCGA-EK-A2H0 | Cisplatin | 84691 | DB00515 | Complete Response | 1 |
| TCGA-C5-A1BQ | Cisplatin | 84691 | DB00515 | Complete Response | 1 |
| TCGA-HG-A2PA | Paclitaxel | 36314 | DB01229 | Partial Response | 0 |
| TCGA-HG-A2PA | Carboplatin | 426756 | DB00958 | Partial Response | 0 |
| TCGA-IR-A3L7 | Cisplatin | 84691 | DB00515 | Complete Response | 1 |
| TCGA-IR-A3LC | Cisplatin | 84691 | DB00515 | Complete Response | 1 |
| TCGA-IR-A3LH | Cisplatin | 84691 | DB00515 | Complete Response | 1 |
| TCGA-IR-A3LI | Cisplatin | 84691 | DB00515 | Complete Response | 1 |
| TCGA-IR-A3LI | Paclitaxel | 36314 | DB01229 | Complete Response | 1 |
| TCGA-IR-A3LI | Topotecan | 60700 | DB01030 | Complete Response | 1 |
| TCGA-IR-A3LI | Cisplatin | 84691 | DB00515 | Complete Response | 1 |
| TCGA-IR-A3LK | Cisplatin | 84691 | DB00515 | Complete Response | 1 |
| TCGA-IR-A3LK | Carboplatin | 426756 | DB00958 | Clinical Progressive Disease | 0 |
| TCGA-IR-A3LK | Paclitaxel | 36314 | DB01229 | Clinical Progressive Disease | 0 |
| TCGA-IR-A3LL | Cisplatin | 84691 | DB00515 | Complete Response | 1 |
| TCGA-EX-A3L1 | Cisplatin | 84691 | DB00515 | Complete Response | 1 |
| TCGA-EA-A3QD | Cisplatin | 84691 | DB00515 | Complete Response | 1 |
| TCGA-IR-A3LA | Cisplatin | 84691 | DB00515 | Complete Response | 1 |
| TCGA-IR-A3LA | Fluorouracil | 3385 | DB00544 | Complete Response | 1 |
| TCGA-EX-A449 | Cisplatin | 84691 | DB00515 | Complete Response | 1 |
| TCGA-EX-A449 | Carboplatin | 426756 | DB00958 | Complete Response | 1 |
| TCGA-EX-A449 | Paclitaxel | 36314 | DB01229 | Complete Response | 1 |
| TCGA-IR-A3LB | Cisplatin | 84691 | DB00515 | Complete Response | 1 |
| TCGA-IR-A3LB | Fluorouracil | 3385 | DB00544 | Complete Response | 1 |
| TCGA-IR-A3LB | Carboplatin | 426756 | DB00958 | Stable Disease | 0 |
| TCGA-IR-A3LB | Paclitaxel | 36314 | DB01229 | Stable Disease | 0 |
| TCGA-LP-A4AU | Cisplatin | 84691 | DB00515 | Complete Response | 1 |
| TCGA-LP-A4AU | Fluorouracil | 3385 | DB00544 | Complete Response | 1 |

| | | | | | |
|---|---|---|---|---|---|
| TCGA-EA-A44S | Carboplatin | 426756 | DB00958 | Stable Disease | 0 |
| TCGA-EA-A44S | Bleomycin | 5460769 | DB00290 | Stable Disease | 0 |
| TCGA-EA-A44S | Gemcitabine | 60750 | DB00441 | Stable Disease | 0 |
| TCGA-EA-A44S | Doxorubicin | 31703 | DB00997 | Stable Disease | 0 |
| TCGA-EA-A44S | Bleomycin | 5460769 | DB00290 | Stable Disease | 0 |
| TCGA-EA-A4BA | Carboplatin | 426756 | DB00958 | Complete Response | 1 |
| TCGA-EA-A4BA | Cyclophosphamide | 2907 | DB00531 | Complete Response | 1 |
| TCGA-EA-A4BA | Doxorubicin | 31703 | DB00997 | Complete Response | 1 |
| TCGA-HM-A4S6 | Cisplatin | 84691 | DB00515 | Complete Response | 1 |
| TCGA-MY-A5BD | Cisplatin | 84691 | DB00515 | Complete Response | 1 |
| TCGA-MY-A5BF | Cisplatin | 84691 | DB00515 | Complete Response | 1 |
| TCGA-DS-A5RQ | Cisplatin | 84691 | DB00515 | Complete Response | 1 |
| TCGA-JW-A5VI | Topotecan | 60700 | DB01030 | Clinical Progressive Disease | 0 |
| TCGA-Q1-A5R2 | Cisplatin | 84691 | DB00515 | Partial Response | 0 |
| TCGA-Q1-A5R3 | Cisplatin | 84691 | DB00515 | Partial Response | 0 |
| TCGA-Q1-A73O | Cisplatin | 84691 | DB00515 | Complete Response | 1 |
| TCGA-RA-A741 | Carboplatin | 426756 | DB00958 | Complete Response | 1 |
| TCGA-RA-A741 | Paclitaxel | 36314 | DB01229 | Complete Response | 1 |
| TCGA-DS-A7WF | Cisplatin | 84691 | DB00515 | Partial Response | 0 |
| TCGA-DS-A7WH | Cisplatin | 84691 | DB00515 | Complete Response | 1 |
| TCGA-DS-A7WI | Cisplatin | 84691 | DB00515 | Complete Response | 1 |

**Supplementary Table 6**. The predicted $IC_{50}$ values of missing data in the GDSC database. Note that only the top-10 drugs with highest efficacy and the corresponding top-10 associated cancer cell lines were shown.

| PubChem id | Drug name | depMapID | Cell line | TCGA type | Predicted ln(IC50) |
|---|---|---|---|---|---|
| 387447 | Bortezomib | ACH-000718 | NCI-H2291 | LUAD | -1.602 |
| 387447 | Bortezomib | ACH-000818 | BT-483 | BRCA | -1.995 |
| 387447 | Bortezomib | ACH-000727 | NCI-H2066 | SCLC | -2.512 |
| 387447 | Bortezomib | ACH-000987 | MeWo | SKCM | -2.835 |
| 387447 | Bortezomib | ACH-000443 | OVKATE | OV | -2.88 |
| 387447 | Bortezomib | ACH-000852 | NCI-H1435 | LUAD | -3.08 |
| 387447 | Bortezomib | ACH-000360 | NCI-H508 | COAD/READ | -3.085 |
| 387447 | Bortezomib | ACH-000680 | SW948 | COAD/READ | -3.457 |
| 387447 | Bortezomib | ACH-000698 | DMS 53 | SCLC | -3.481 |
| 387447 | Bortezomib | ACH-000489 | SW1116 | COAD/READ | -3.529 |
| 148124 | Docetaxel | ACH-000727 | NCI-H2066 | SCLC | -1.53 |
| 148124 | Docetaxel | ACH-000568 | UACC-812 | BRCA | -2.351 |
| 148124 | Docetaxel | ACH-000752 | NCI-H196 | SCLC | -2.862 |
| 148124 | Docetaxel | ACH-000999 | SNU-1040 | COAD/READ | -2.863 |
| 148124 | Docetaxel | ACH-000386 | KG-1 | LAML | -2.976 |
| 148124 | Docetaxel | ACH-000489 | SW1116 | COAD/READ | -2.981 |
| 148124 | Docetaxel | ACH-000470 | SW1463 | COAD/READ | -3.042 |
| 148124 | Docetaxel | ACH-000197 | TALL-1 | ALL | -3.23 |
| 148124 | Docetaxel | ACH-000399 | NCI-H2196 | SCLC | -3.309 |
| 148124 | Docetaxel | ACH-000373 | SKM-1 | LAML | -3.371 |
| 448013 | Epothilone B | ACH-000727 | NCI-H2066 | SCLC | -0.188 |
| 448013 | Epothilone B | ACH-000828 | ZR-75-30 | BRCA | -1.309 |
| 448013 | Epothilone B | ACH-000489 | SW1116 | COAD/READ | -1.562 |
| 448013 | Epothilone B | ACH-000347 | QGP-1 | PAAD | -1.636 |
| 448013 | Epothilone B | ACH-000360 | NCI-H508 | COAD/READ | -2.005 |
| 448013 | Epothilone B | ACH-000798 | CL-40 | COAD/READ | -2.041 |
| 448013 | Epothilone B | ACH-000626 | U266B1 | MM | -2.124 |
| 448013 | Epothilone B | ACH-000640 | SK-MEL-31 | SKCM | -2.42 |
| 448013 | Epothilone B | ACH-000678 | MKN7 | STAD | -2.552 |
| 448013 | Epothilone B | ACH-000373 | SKM-1 | LAML | -2.758 |
| 6710780 | Vinblastine | ACH-000727 | NCI-H2066 | SCLC | -0.237 |
| 6710780 | Vinblastine | ACH-000568 | UACC-812 | BRCA | -1.649 |
| 6710780 | Vinblastine | ACH-000470 | SW1463 | COAD/READ | -1.713 |
| 6710780 | Vinblastine | ACH-000752 | NCI-H196 | SCLC | -1.854 |

| | | | | | |
|---|---|---|---|---|---|
| 6710780 | Vinblastine | ACH-000489 | SW1116 | COAD/READ | -1.937 |
| 6710780 | Vinblastine | ACH-000999 | SNU-1040 | COAD/READ | -2.068 |
| 6710780 | Vinblastine | ACH-000345 | KP-N-RT-BM-1 | NB | -2.484 |
| 6710780 | Vinblastine | ACH-000422 | SNU-475 | LIHC | -2.535 |
| 6710780 | Vinblastine | ACH-000035 | NCI-H1650 | LUAD | -2.596 |
| 6710780 | Vinblastine | ACH-000615 | SK-MEL-28 | SKCM | -2.664 |
| 11178236 | Sepantronium bromide | ACH-000727 | NCI-H2066 | SCLC | 0.435 |
| 11178236 | Sepantronium bromide | ACH-000798 | CL-40 | COAD/READ | 0.193 |
| 11178236 | Sepantronium bromide | ACH-000103 | Caov-4 | OV | -1.089 |
| 11178236 | Sepantronium bromide | ACH-000429 | A-704 | KIRC | -1.132 |
| 11178236 | Sepantronium bromide | ACH-000828 | ZR-75-30 | BRCA | -1.187 |
| 11178236 | Sepantronium bromide | ACH-000360 | NCI-H508 | COAD/READ | -1.358 |
| 11178236 | Sepantronium bromide | ACH-000626 | U266B1 | MM | -1.492 |
| 11178236 | Sepantronium bromide | ACH-000895 | CL-34 | COAD/READ | -1.522 |
| 11178236 | Sepantronium bromide | ACH-000640 | SK-MEL-31 | SKCM | -1.617 |
| 11178236 | Sepantronium bromide | ACH-000822 | SK-MEL-24 | SKCM | -2.349 |
| 36314 | Paclitaxel | ACH-000818 | BT-483 | BRCA | 1.265 |
| 36314 | Paclitaxel | ACH-000443 | OVKATE | OV | 1.002 |
| 36314 | Paclitaxel | ACH-000727 | NCI-H2066 | SCLC | 0.686 |
| 36314 | Paclitaxel | ACH-000680 | SW948 | COAD/READ | 0.215 |
| 36314 | Paclitaxel | ACH-000347 | QGP-1 | PAAD | 0.058 |
| 36314 | Paclitaxel | ACH-000718 | NCI-H2291 | LUAD | -0.023 |
| 36314 | Paclitaxel | ACH-000828 | ZR-75-30 | BRCA | -0.099 |
| 36314 | Paclitaxel | ACH-000987 | MeWo | SKCM | -0.147 |
| 36314 | Paclitaxel | ACH-000568 | UACC-812 | BRCA | -0.319 |
| 36314 | Paclitaxel | ACH-000986 | HT115 | COAD/READ | -0.522 |
| 446378 | Thapsigargin | ACH-000727 | NCI-H2066 | SCLC | 1.129 |
| 446378 | Thapsigargin | ACH-000798 | CL-40 | COAD/READ | 0.054 |
| 446378 | Thapsigargin | ACH-000828 | ZR-75-30 | BRCA | 0.002 |
| 446378 | Thapsigargin | ACH-000347 | QGP-1 | PAAD | -0.126 |

| | | | | | |
|---|---|---|---|---|---|
| 446378 | Thapsigargin | ACH-000360 | NCI-H508 | COAD/READ | -0.594 |
| 446378 | Thapsigargin | ACH-000489 | SW1116 | COAD/READ | -0.744 |
| 446378 | Thapsigargin | ACH-000895 | CL-34 | COAD/READ | -0.86 |
| 446378 | Thapsigargin | ACH-000148 | Hs 578T | BRCA | -0.948 |
| 446378 | Thapsigargin | ACH-000626 | U266B1 | MM | -1.149 |
| 446378 | Thapsigargin | ACH-000678 | MKN7 | STAD | -1.473 |
| 5311497 | Vinorelbine | ACH-000727 | NCI-H2066 | SCLC | 0.287 |
| 5311497 | Vinorelbine | ACH-000347 | QGP-1 | PAAD | -0.456 |
| 5311497 | Vinorelbine | ACH-000489 | SW1116 | COAD/READ | -1.054 |
| 5311497 | Vinorelbine | ACH-000828 | ZR-75-30 | BRCA | -1.124 |
| 5311497 | Vinorelbine | ACH-000798 | CL-40 | COAD/READ | -1.452 |
| 5311497 | Vinorelbine | ACH-000360 | NCI-H508 | COAD/READ | -1.492 |
| 5311497 | Vinorelbine | ACH-000895 | CL-34 | COAD/READ | -1.579 |
| 5311497 | Vinorelbine | ACH-000999 | SNU-1040 | COAD/READ | -1.811 |
| 5311497 | Vinorelbine | ACH-000532 | SNU-61 | COAD/READ | -1.814 |
| 5311497 | Vinorelbine | ACH-000725 | HCC202 | BRCA | -1.948 |
| 104842 | SN-38 | ACH-000568 | UACC-812 | BRCA | -1.543 |
| 104842 | SN-38 | ACH-000399 | NCI-H2196 | SCLC | -2.393 |
| 104842 | SN-38 | ACH-000953 | SUP-T1 | ALL | -2.428 |
| 104842 | SN-38 | ACH-000386 | KG-1 | LAML | -2.524 |
| 104842 | SN-38 | ACH-000345 | KP-N-RT-BM-1 | NB | -2.587 |
| 104842 | SN-38 | ACH-000197 | TALL-1 | ALL | -2.807 |
| 104842 | SN-38 | ACH-000422 | SNU-475 | LIHC | -3.006 |
| 104842 | SN-38 | ACH-000713 | Caov-3 | OV | -3.394 |
| 104842 | SN-38 | ACH-000515 | HCC-33 | SCLC | -3.488 |
| 104842 | SN-38 | ACH-000525 | NCI-H2171 | SCLC | -3.663 |
| 25167777 | Omipalisib | ACH-000727 | NCI-H2066 | SCLC | 1.437 |
| 25167777 | Omipalisib | ACH-000626 | U266B1 | MM | 0.502 |
| 25167777 | Omipalisib | ACH-000640 | SK-MEL-31 | SKCM | -1.145 |
| 25167777 | Omipalisib | ACH-000798 | CL-40 | COAD/READ | -1.173 |
| 25167777 | Omipalisib | ACH-000828 | ZR-75-30 | BRCA | -1.791 |
| 25167777 | Omipalisib | ACH-000288 | BT-549 | BRCA | -1.823 |
| 25167777 | Omipalisib | ACH-000373 | SKM-1 | LAML | -1.914 |
| 25167777 | Omipalisib | ACH-000895 | CL-34 | COAD/READ | -1.915 |
| 25167777 | Omipalisib | ACH-000360 | NCI-H508 | COAD/READ | -2.105 |
| 25167777 | Omipalisib | ACH-000858 | KNS-62 | LUSC | -2.469 |

**Supplementary Table 7**. We have applied both binomial exact test and Mann-Whitney U test to verify the superiority of our method. As for the former, we count the number of drugs that our method achieves a larger PCC than a baseline method and then test with the alternative hypothesis that the probability that our method outperforms the baseline is greater than 0.5. As for the later, we test with the alternative hypothesis that the PCCs of our method for the different drugs have a positive shift when compared with those of a baseline.

| | DeepCDR vs tCNNs |
|---|---|
| Exact binomial test | $2.2\times10^{-16}$ |
| Mann-Whitney U test | $1.51\times10^{-06}$ |

**Supplementary Table 8**. We have applied both binomial exact test and Mann-Whitney U test to verify the superiority of our method. As for the former, we count the number of cell lines that our method achieves a larger PCC than a baseline method and then test with the alternative hypothesis that the probability that our method outperforms the baseline is greater than 0.5. As for the later, we test with the alternative hypothesis that the PCCs of our method for the different cell lines have a positive shift when compared with those of a baseline.

| | DeepCDR vs tCNNs |
|---|---|
| Exact binomial test | $2.2\times10^{-16}$ |
| Mann-Whitney U test | $1.06\times10^{-22}$ |

**Supplementary Table 9**. We have applied both binomial exact test and Mann-Whitney U test to verify the superiority of our method. As for the former, we count the number of drugs that our method achieves a larger AUC than a baseline method and then test with the alternative hypothesis that the probability that our method outperforms the baseline is greater than 0.5. As for the later, we test with the alternative hypothesis that the AUCs of our method for the different drugs have a positive shift when compared with those of a baseline.

| | DeepCDR vs tCNNs |
|---|---|
| Exact binomial test | $2.2\times10^{-16}$ |
| Mann-Whitney U test | $4.23\times10^{-5}$ |

**Supplementary Table 10**. We have applied both binomial exact test and Mann-Whitney U test to verify the superiority of our method. As for the former, we count the number of cell lines that our method achieves a larger AUC than a baseline method and then test with the alternative hypothesis that the probability that our method outperforms the baseline is greater than 0.5. As for the later, we test with the alternative hypothesis that the AUCs of our method for the different cell lines have a positive shift when compared with those of a baseline.

| | DeepCDR vs tCNNs |
|---|---|
| Exact binomial test | $2.2\times10^{-16}$ |
| Mann-Whitney U test | $1.39\times10^{-35}$ |
